# Sequence Controlled Secondary Structure Determines Site-selectivity of Lanthipeptides

**DOI:** 10.1101/2022.11.28.518241

**Authors:** Xuenan Mi, Emily K. Desormeaux, Tung T. Le, Wilfred A. van der Donk, Diwakar Shukla

## Abstract

Lanthipeptides are ribosomally synthesized and post-translationally modified peptides that are generated from precursor peptides through a dehydration and cyclization process in the biosynthetic pathways. In contrast to most other lanthipeptide synthetases, ProcM, a class II lanthipeptide synthetase, demonstrates high substrate tolerance. It is enigmatic that a single enzyme can catalyze the cyclization process of a diverse range of substrates with high fidelity. Previous studies suggested that the site-selectivity of lanthionine formation is determined by substrate sequence rather than by the enzyme. However, exactly how substrate sequence contributes to site-selective lanthipeptide biosynthesis is not clear. In this study, we performed molecular dynamic simulations for ProcA3.3 core peptide variants to explore how the predicted solution structure of the substrate without enzyme correlates to final product formation. Our simulation results support a model in which the secondary structure of the core peptide controls the ring pattern of the final product. We also demonstrate that the dehydration step in the biosynthesis pathway does not influence the site-selectivity of ring formation. In addition, we performed simulation for the core peptides of ProcA1.1 and 2.8, which are well-suited candidates to investigate the connection between order of ring formation and solution structure. Simulation results indicate that in both cases, C-terminal ring formation is more likely which was supported by experimental results. Our findings indicate that the substrate sequence and its solution structure can be used to predict the site-selectivity and order of ring formation, and that secondary structure is a crucial factor influencing the site-selectivity. Taken together, these findings will facilitate our understanding of the lanthipeptide biosynthetic mechanism and accelerate bioengineering efforts for lanthipeptide-derived products.

## Introduction

Natural products are important sources of new drugs, with more than 60% of all new drugs derived from natural products or their derivatives.^1^ Ribosomally synthesized and post-translationally modified peptides (RiPPs) are a fast-growing natural product family because recent developments in genome mining algorithms have facilitated their discovery. ^2–8^ Most RiPPs follow a similar biosynthetic logic: the precursor peptide, encoded by a structural gene, is modified by enzymes to generate the mature natural product. Most precursor peptides are composed of a highly conserved leader peptide at the N-terminus, which is important for recognition by modification enzymes, as well as a highly variable core sequence at the C-terminus that is transformed into the mature product.^9^

Lanthipeptides are the largest group of RiPPs based on currently sequenced genomes^10,11^ and have a variety of bioactivities, including antimicrobial,^12^ antiviral,^13^ antifungal^14^ and antiallodynic activities.^15^ There are multiple classes of enzyme that are able to install lanthionine (Lan) or methyllanthionine (MeLan) rings. This paper will discuss class II lanthipeptide synthetases which are single bifunctional enzymes capable of catalyzing both the dehydration and the cyclization reactions. These lanthipeptide synthetases catalyze the dehydration of Ser/Thr residues to dehydroalanine (Dha)/ (Z)-dehydrobutyrine (Dhb) respectively, followed by an intramolecular Michael-type addition of Cys thiols to the Dha/Dhb to form the class-defining Lan or MeLan linkages (Figure 1A, B). ^16^ After modification, the leader peptide is removed by proteases and the mature product is exported.

**Figure 1:**
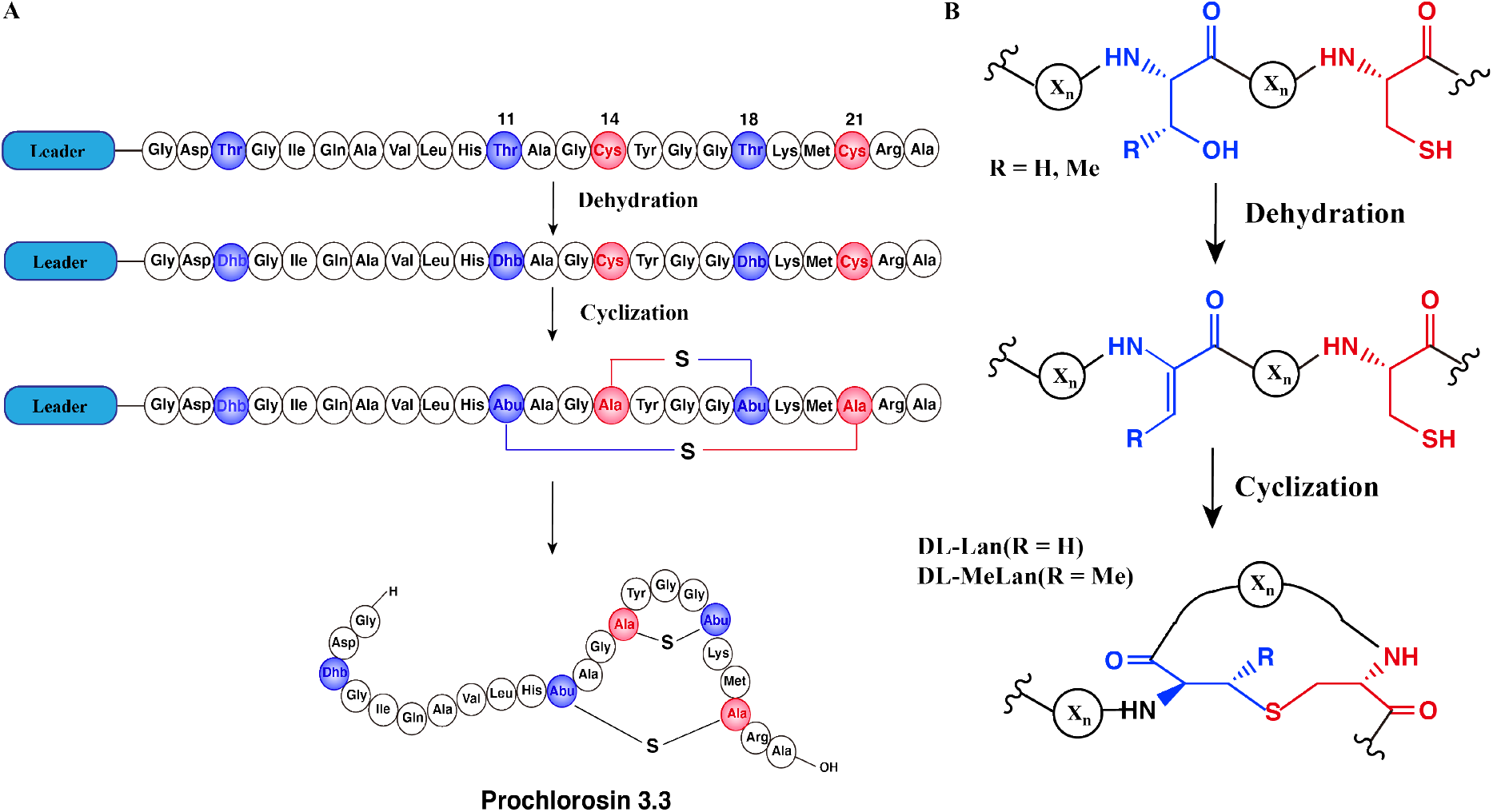
Schematic Biosynthetic Pathway of Prochlorosin 3.3. (A) Biosynthetic route to Pcn3.3 and cartoon representation of the sequence and structure. (B) Post-translational modifications carried out by ProcM during Pcn3.3 biosynthesis. X_*n*_ represents the peptide chain.

Enzymatic modifications catalyzed by many lanthipeptide synthetases demonstrate relaxed substrate specificity, often tolerating significant changes made to the core peptide sequences. ^17–20^ This feature has been widely utilized for bioengineering of novel lanthipeptides.^21–28^ A remarkable example of substrate tolerance is the class II lanthipeptide synthetase ProcM, which was discovered in the marine picocyanobacterium *Prochlorococcus* MIT9313.^29^ Unlike most other lanthipeptide biosynthetic pathways, the ProcM biosynthetic gene cluster does not encode just a single precursor peptide with one modifying enzyme. ^29^ Along with ProcM, genes encoding 30 distinct putative precursor peptides were identified in the genome, termed ProcA; the mature products were called prochlorosins (Pcn). A related enzyme SyncM is encoded in Synechococcus MITS9509 together with up to 80 putative substrate peptides, ^30^ and SyncM has also been demonstrated to have very high substrate tolerance.^31^ ProcM is a bifunctional enzyme with an N-terminal dehydratase domain and a C-terminal cyclase domain that acts on these 30 distinct substrate sequences, and forms lanthipeptides with highly diverse sequences and ring patterns. ^29,32,33^ Previous studies have provided a possible explanation suggesting that the core peptide sequence rather than the enzyme may determine the final ring pattern.^34–36^ In this proposed model, the substrate has a propensity towards a specific ring pattern based on its conformational free energy landscape. ProcM would accelerate the cyclization by increasing the nucleophilicity of the Cys thiolate coordinated to the zinc ion in the active site and covalently locking the peptide into such favorable conformations. ^35^

In a recent study, Le et al. investigated how substrate sequence controls site-selectivity of lanthionine formation by ProcM. ^34^ The findings with a library of ProcA3.3 precursor peptide variants supported a model in which substrate sequence determines the site-selectivity of lanthionine formation. However, the experimental data could not predict how the core sequence controls the ring pattern of the final product through modulating its conformational free energy landscape. Computational studies using molecular dynamics (MD) simulations have been an effective approach to investigate conformational equilibria of peptides and proteins.^37–41^ In this study we used the core peptide of ProcA3.3 as a model and performed atomic-scale molecular dynamic simulations of linear peptides in solution to explore the factors that may determine the ring pattern. Our results revealed that the secondary structure of the core sequence is an important factor for controlling the final product structure. Furthermore, we investigated whether the order of cyclization is determined by substrate sequence alone or is also influenced by ProcM. Pcn1.1 and Pcn2.8, which contain two non-overlapping rings, are well-suited candidates to study the order of cyclization. Our computational analysis indicated that the C-terminal ring of ProcA1.1 and ProcA2.8 have higher probabilities to form based on energetic stabilities inferred from the simulation-based conformational energy landscape. To validate our results, we performed Liquid Chromatography-Mass Spectrometry (LC-MS) analysis to confirm that ProcA1.1 is cyclized in a C-to-N-terminal fashion. Our work provides computational evidence to explain how substrate sequence determines the prochlorosin site-selectivity and order of cyclization and demonstrates how secondary structure of the core sequence is a crucial factor to control ring patterns of final lanthipeptide products.

## Methods

### Molecular Dynamics (MD) simulation

Atomistic MD simulations were performed starting from the linear structure of the peptides. All peptides were constructed using PyMol. ^42^ The peptide was solvated in a TIP3P water box, and the system was neutralized by Na^+^ ions and Cl^−^ ions using Packmol.^43^ All MD simulations were performed using the Amber18 software package employing Amber ff14SB force field.^44^ Parameters for non-natural dehydro amino acids were generously provided by Gonzalo Jiménez-Osés. ^34^ They were generated originally using the AMBER gaff2 force field and with partial charges set to fit the electrostatic potential generated with HF/6-31G(d) using the RESP method.^45^ The charges were calculated according to the Merz-Singh-Kollman scheme using Gaussian 16.^46^ Each MD system was first minimized for 50,000 cycles using steepest descent for the first 5,000 cycles and conjugate gradient for the remaining 45,000 cycles. The systems were heated from 0 K to 300 K under NVT ensemble. The heating step was conducted for 3 ns using Langevin thermostat with a collision frequency of 2 ps^−1^.^47^ The systems were equilibrated in NPT ensemble for 2 ns with a pressure of 1 bar using Monte Carlo barostat.^48^ The systems were further equilibrated in NPT ensemble (300 K and 1 bar) for 50 ns and then underwent production runs.^49^ The SHAKE algorithm was used to constrain hydrogen-containing bonds.^50^ All systems were subject to hydrogen mass repartitioning (HMR),^51^ which redistributes mass between hydrogen atoms and covalently bonded atoms of peptide to allow the time step of the simulation to be increased to 4 fs.

### Adaptive sampling

All simulation data were obtained by applying adaptive sampling method, that was used to efficiently sample the conformational space of the peptide. ^52^ The adaptive sampling approach has been applied in sampling protein-ion binding, protein-ligand binding and protein folding processes.^53–56^ In this study, the least count based adaptive sampling was used to find the new conformational states quickly. ^52^ Adaptive sampling was performed as follows:

1. Run a series of short MD simulations from a collection of starting structures.
2. Cluster all collected simulation data using a K-means algorithm. ^57^ The distances of all pairs of residues separated by two or more residues were used as features to generate 100 clusters.
3. Randomly pick one state from each of 25 clusters with the least population as seeds to start new simulation.
4. Repeat steps 1-3 until the sampling reaches convergence.

### Markov State Model (MSM) construction

In the adaptive sampling, we generated many short trajectories to capture the dynamic process of the system. MSM was built to connect these independent trajectories thermodynamically and kinetically, and remove bias introduced by the least count based adaptive sampling. ^54,58,59^ The distances between residue pairs separated by two or more residues were used to featurize the simulation data. The time-lagged component analysis (tICA) was used to reduce dimensions of featurized data. Each time-lagged component (tIC) is a linear combination of features. Using tICs, we could determine the discrete states of the system, which have different slowest timescales. The discrete states were further discretized based on K-means algorithm. ^57^ Then MSM is used to model the entire dynamic system through the corresponding transition probability matrix (*T*) between discretized states. Each element (*T*_*ij*_) in the matrix represents the probability of transitioning from state *i* to state *j* at lag time *τ*, which is long enough to be validate Markovian behavior. ^59^ The lag time *τ* is estimated from the implied time scale, the minimum *τ* at which the implied time scale converged was selected as the MSM lag time (Figure S7-S13A). The optimized hyperparameters (the number of clusters and the number of tICs) of the MSM were selected by maximizing the VAMP-2 score (Figure S7-S13B), calculated by the sum of squared eigenvalues from the transition probability matrix,^60,61^ which maximizes the kinetic variance contained in the features. ^62,63^ The first six eigenvalue were used to calculate the VAMP-2 score and 10-fold cross-validation was calculated to obtain the average score using pyEMMA. ^64^

### Trajectory analysis

CPPTRAJ module in AMBER 18^44^ and the python package MDTraj 1.9.0^65^ were used to analyze trajectory data, and VMD 1.9.3^66^ and PyMol^42^ were used to visualize MD snapshots. The python package Matplotlib^67^ was used to generate the 2-D plot of the free energy landscape.

### Ring formation probability calculation

The ring formation was defined as the distance between the *β*-carbon of threonine and the sulfur of cysteine being within 7.5 Å. The distance between atoms was calculated using MDtraj 1.9.0. ^65^ The equilibrium ring formation probability was calculated as the product of raw ring formation probability within each MSM state multiplied by the equilibrium probability of the MSM state, as follows:

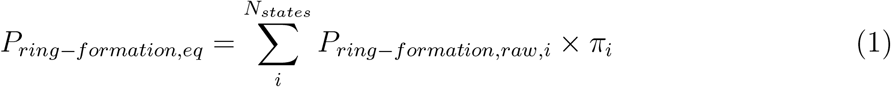

### Error analysis

Errors on thermodynamics calculations were generated by bootstrapping. ^68^ For each bootstrap sample, 80% of total number of trajectories were randomly selected. We kept the original state index and built MSM for each sample. We generated N bootstrap samples and used the standard deviation of N samples as error bar.

### Purification of ProcA peptides

The genes for His_6_-tagged ProcA peptides were cloned into a pET15b vector and overexpressed as previously described.^69^ Cell paste was resuspended in 20mL of LanA B1 buffer (6.0 M guanidine HCl, 20 mM NaH_2_PO_4_, 0.5 mM imidazole, 1 mM TCEP, pH 7.5) for 1 L of cell culture and lysed via sonication at 60% amplitude for 5 min with a 2.0 s on and 6.0 s off pulse. The lysate was clarified via centrifugation at 50,000xg for 1 h and the supernatant was filtered using 0.45 *μ*m syringe filters. The filtered lysate was loaded onto a gravity flow Ni-NTA column with 1mL of resin preequilibrated with 6 column volumes (CV) of LanA B1 buffer. The column was washed with 10 CV of LanA B1 buffer, 5 CV of LanA B2 buffer (6.0 M guanidine HCl, 20 mM NaH_2_PO_4_, 30 mM imidazole, 1 mM TCEP, pH 7.5), and 5 CV of LanA wash buffer (20 mM NaH_2_PO_4_, 30 mM imidazole, 300 mM NaCl, 1 mM TCEP, pH 7.5). The peptide was then eluted using 10 CV of LanA elution buffer (20 mM NaH_2_PO_4_, 1 M imidazole, 100 mM NaCl, 1 mM TCEP, pH 7.5). The buffer of the purified peptide was then exchanged into a thrombin digestion buffer (50 mM HEPES, 100 mM NaCl, 1 mM TCEP, pH 8.0) using a 3 kDa Amicon centrifugation filter and the sample was concentrated to approximately 5 mL. Bovine Thrombin High Purify Grade (MP Biomedicals) was added to the sample (100 units) and the sample was left overnight at 4*°C* for removal of the His_6_-tag. Thrombin-digested peptides were purified via reversed phase HPLC using a Phenomenex Luna C5 semiprep column (250×10 mm, 10 *μ*, 100 Å) at a flow rate of 8 mL/min and with the following gradient over 32 min: 2% B for 10 min, 2-30% B over 2 min, 30-60% B over 15 min, 60-100% B over 2 min, and hold at 100% B for 3 min (A: 0.1% TFA in H_2_O, B:100% MeCN, 0.1% TFA). HPLC-purified peptide was collected and lyophilized, and the peptides were stored as a powder at −20*°C* until use.

### Purification of ProcM

ProcM was overexpressed as reported earlier utilizing a pRSFDuet vector. ^69,70^ All purification steps were carried out in a cold room (4*°C*) or on an ice bath. Cell paste was resuspended in 20 mL of LanM start buffer (20 mM Tris, 500 mM NaCl, 10% glycerol, pH 7.6) per L of expression and the cell mixture was allowed to nutate with protease inhibitor (Pierce^*T M*^), lysozyme (50 mg/L culture), and benzonase (12 *μ*L/L cell culture) for 1 h. Cell lysis was performed via sonication at 35% amplitude for 15 min with a 4.0 s on, 9.9 s off pulse. The lysate was clarified via centrifugation at 50,000xg for 1 h and the supernatant was filtered using 0.45 *μ*m syringe filters. The filtered lysate was loaded onto a Ni-Hi-Trap column equilibrated with 6 CV of ProcM start buffer. The column was manually washed by 6 CV of ProcM wash buffer (20 mM Tris, 500 mM NaCl, 30 mM imidazole, 10% glycerol, pH 7.6) before being attached to an Akta fast protein liquid chromatography (FPLC) system to complete the elution using ProcM wash and elution buffers (20 mM Tris, 500 mM NaCl, 1 M imidazole, 10% glycerol, pH 7.6) and monitoring elution at 280 nm with the following gradient: 0-2% over 2 CV, 2-20% over 10 CV, 20-100% over 0.5 CV. The protein elution fractions were analyzed via gel electrophoresis and fractions containing ProcM were concentrated using 50 kDa Amicon centrifugal filtration. The ProcM was then further purified/desalted using a FPLC gel filtration column (Superdex 200, 1.5 × 60 cm, GE healthcare) using a 1 mL/min flow rate and an isocratic elution of ProcM storage buffer (50 mM HEPES, 500 mM KCl, 5% glycerol, pH 7.6). ProcM eluted in three peaks (aggregate, then oligomer, followed by monomer). The monomeric peaks were collected, concentrated via 50 kDa Amicon centrifugal filtration, aliquoted into single use portions, flash frozen in liquid nitrogen, and stored at −80*°C* until use.

### ProcM reactions

Prior to initiation of reactions, 4 *μ*M ProcM and 80 *μ*M ProcA peptide were preincubated separately at 25*°C* for 1 h in 750 *μ*L of reaction mixture (5 mM ATP, 0.17 mM ADP, 5 mM MgCl_2_, 100 mM HEPES, 0.1 mM TCEP, pH 7.5). The two samples were mixed thoroughly to initiate a reaction with a final concentration of 2 *μ*M ProcM and 40 *μ*M ProcA peptide. An 80 *μ*L aliquot was removed at desired time points and quenched into 900 *μ*L of quench buffer (111 mM citrate, 1.11 mM EDTA, pH 3.0). Each reaction was initially quenched at 15, 30, 45, and 60 min. Reaction times were adjusted as needed to obtain monocyclized intermediate (30 min for ProcA1.1). After quenching, 100 *μ*L of 100 mM TCEP was added to each aliquot and samples were incubated at 25*°C* for 10 min. The pH of the aliquots was adjusted to approximately 6.3 via the addition of 40-45 *μ*L of 5 M NaOH. The free thiols were then alkylated by the addition of 11 *μ*L of 1 M *N* -ethylmaleimide (NEM) in EtOH and the reaction was incubated at 37*°C* for 10 min before the reaction was quenched by the addition of 11 *μ*L of formic acid. ^71^ The buffer of the time points was then exchanged to water using 3 kDa Amicon centrifugal filters. Samples were then lyophilized and stored as a powder until used for analysis.

### Liquid Chromatography-Mass Spectrometry

Samples were resuspended in 50 *μ*L of water, and the leader peptide was removed via the addition of 50 *μ*L of LahT_150_ solution (2.0 mg/mL) and left overnight. The protease LahT_150_ ^72^ was removed from samples via addition of 100 *μ*L of MeOH and 5 *μ*L of formic acid. The sample was centrifuged to pellet any precipitated protein in the sample and 10 *μ*L aliquots were injected onto an Agilent Infinity1260 Liquid Chromatography instrument coupled to an Agilent 6454 QTOF mass spectrometer using a Kinetex® 2.6 *μ*m C8 100 Å, LC column (150 × 2.1 mm). The samples were run with a flow rate of 0.4 mL/min at 45 °C. The peptides were eluted using the following gradient: 0-5% B over 3 min, 5-50% B over 10 min, and 50-95% B over 1 min for a total run time of 14 min (A: 0.1% formic acid in water, B: 100% MeCN, 0.1% formic acid). QTOF data was collected with the following settings: polarity= positive, data storage= centroid, acquisition range (m/z)= 100-1700, gas temperature= 325*°C*, gas flow= 13 L/min, nebulizer= 35 psig, sheath gas temp= 275*°C*, sheath gas flow= 12 L/min, Vcap= 4000, nozzle voltage= 500 V, fragmentor=175 V, skimmer= 65 V, octupole RF peak= 750, scan rate= 5 spectra/s, acquisition mode= target MS, and collision energy= 20*/*30 V. Fragmentation at fixed collision energy (20*/*30 V) was performed on the +2*/* + 3 charge species of the target precursors. The data were processed with Qualitative Analysis 10.0 (Agilent).

## Results and Discussion

### Secondary structure may determine ring patterns of ProcA3.3 variants

Previous studies suggested that it is not the enzyme ProcM but rather its substrate sequences that determine the site-selectivity of lanthionine formation. ^34,35^ Recent work applied trapped ion mobility spectrometry-tandem mass spectrometry (TIMS-MS/MS) to discern that wild type (WT) ProcA3.3 contains an overlapping ring pattern, whereas ProcA3.3 Variant1 displays a non-overlapping ring pattern and ProcA3.3 Variant2 displays a mixture of both ring patterns (the ratio of overlapping and non-overlapping is 40:60) (sequences are shown in Figure 2). However, experimental results could not predict how the core sequence controls the ring pattern of the final product. We hypothesized that the solution structure of the core peptide may be a key factor to control ring pattern. ProcM processing of ProcA3.3 has been shown previously to be an irreversible process under assay conditions, and thus the first ring that is formed determines the ultimate ring pattern.^73^

**Figure 2:**
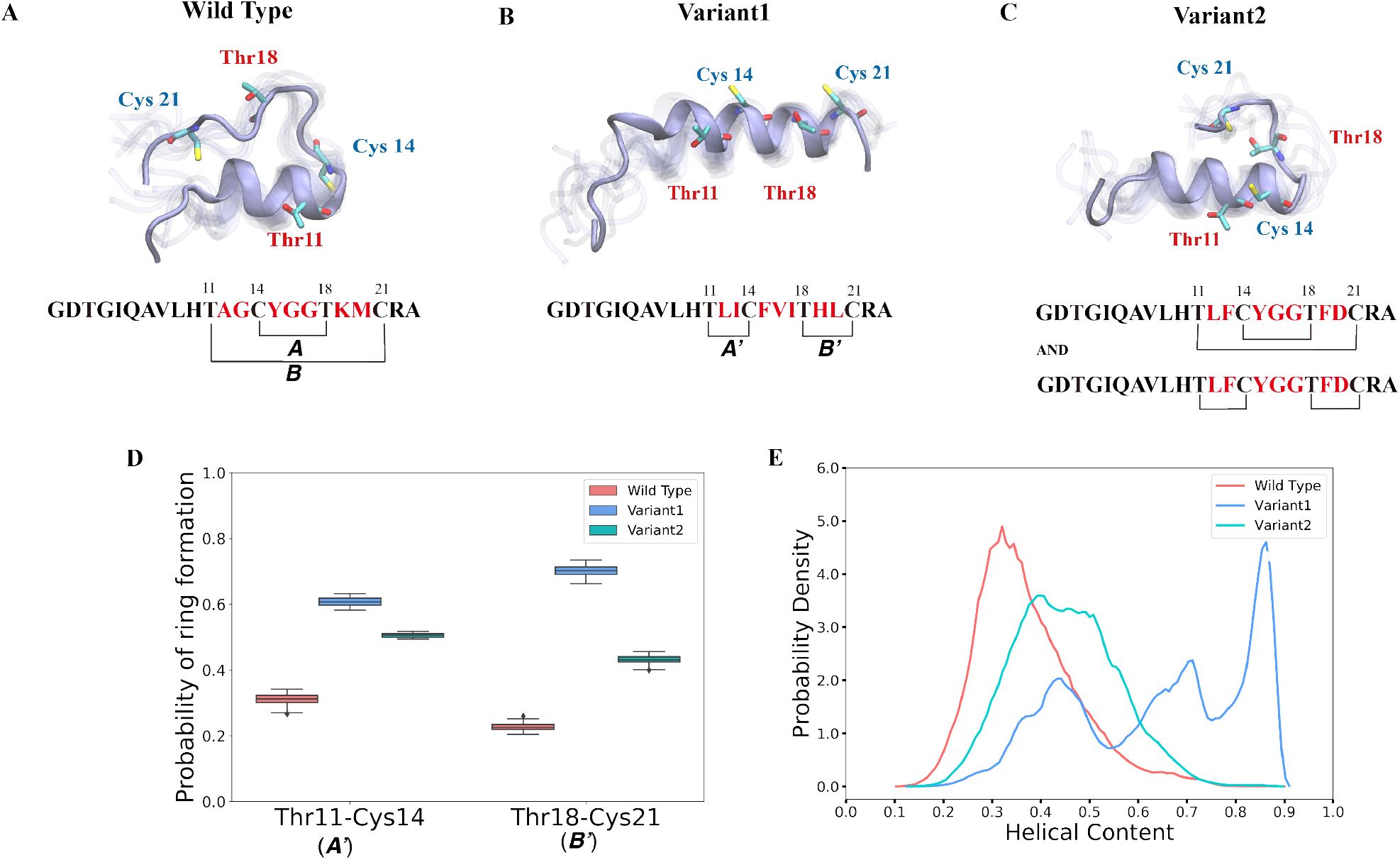
Different secondary structures lead to distinct ring patterns of ProcA3.3 variants. (A, B, C) Cartoon representation of the most populated state in the free energy landscape of ProcA3.3 variants (for the entire free energy landscape of all states, see Figure S2A, B, C). In each state, 10 snapshots were randomly selected and aligned together. One snapshot is shown in ice blue, while the others are shown in transparent ice blue. (D) Probability of formation of ring A′ (Thr11/Cys14) and B′ (Thr18/Cys21) for ProcA3.3 variants. Error bar was calculated by 100 bootstrap samples with 80% of total trajectories. (E) Distribution of helical content of ProcA3.3 variants.

In our study, we first ran 40*μ*s MD simulations for the core peptide of WT ProcA3.3 and these two representative variants to provide the general energy landscape and identify any secondary structures. To evaluate the important residue-residue contacts, we performed time-independent components analysis (tICA) on the simulation data. tICA is a method to identify the slow process by finding coordinates of maximal autocorrelation at a given lag time.^62^ Each time-independent component (tIC) is a linear combination of different features. The first two tICs are the slowest components, and Figure S1 shows that the distance between Thr11 and Cys14 has a high correlation with tIC1, especially for Variant1, which suggests that the contact between Thr11 and Cys14 is one of the slowest dynamic processes during lanthionine formation. We projected the MD simulation data along Thr11-Cys14 distance and *α*−helix content of ProcA3.3 to compare ProcA3.3 WT and variants. Figure 2A shows the most populated conformation from the MSM weighted free energy landscape (Figure S2A) for WT ProcA3.3. We observed that WT ProcA3.3 forms an *α*-helix spanning residues 4 to 12, with Cys14, Thr18 and Cys21 all located within a flexible loop region. This structure is conducive for formation of rings A (Cys14/Thr18) and B (Thr11/Cys21). Conversely, the MSM weighted free energy landscape (Figure S2B) for ProcA3.3 Variant1 shows that the *α*−helix spans residues 8 to 22 (Figure 2B). Thr11, Cys14, Thr18 and Cys21 are all located within the helix, which renders the Thr11-Cys14 residue pair closer and facilitates formation of rings A′ (Thr11/Cys14) and B′ (Thr18/Cys21). For ProcA3.3 Variant2, Thr11 and Cys14 are both within the *α*−helix, but Thr18 and Cys21 are in the flexible loop region (Figure 2C). The *α*−helix structure enables ring A′ and B′ formation; simultaneously, there is a chance to form ring A and B. Therefore, ProcA3.3 Variant2 can form products with both overlapping and non-overlapping ring patterns.

We next compared probabilities of ring formation quantitatively. The ring formation was defined as conformers in which the distance between the *β*-carbon of Thr and the sulfur of Cys was within 7.5 Å. When comparing the probabilities of forming rings A′ and B′ in ProcA3.3 WT and variants, the two variants clearly have a much higher probability than the WT (Figure 2D). Because the A and B rings have different ring sizes compared to ring A′ and B′, preventing direct comparisons, we compared the probabilities of formation of these two sets of rings separately. When comparing the probabilities of forming the ring A, Variant1 has a higher probability of forming this ring than WT (Figure S3A), because *α*−helix also facilitate Cys14 and Thr18 to be close. However, the key question is whether the A′ or B′ rings of Variant1 would form even faster than ring A, which we currently cannot determine from the simulations. We do note that WT has a relatively higher probability of forming ring B (Figure S3B). Here, the distance between the *β*-carbon of Thr and the sulfur of Cys was used as a proxy for the likelihood of ring formation, because the ring is formed between these two atoms. A ring can also be defined to have formed between two residues if the distance between any of their heavy atoms is within a cutoff of 4.5 Å. Using this definition, we observed the same trend of probabilities of different rings formation among variants of the ProcA3.3 core peptide (Figure S3C, S3D, S4). In addition, we quantitatively compared the *α*-helix content in WT and variant ProcA3.3 core peptide. Figure 2E shows that Variant1 has a significantly higher *α*-helix content compared to WT and Variant2.

### Dehydration does not influence ring pattern predictions

Our computational analysis supports the model that substrate sequence determines the cyclization process of ProcA and suggests this occurs at least in part through controlling secondary structure. As mentioned, ProcM processing of ProcA3.3 is an irreversible process under assay conditions, and therefore the first ring that is formed determines the ultimate ring pattern.^73^ In the biosynthesis pathway of lanthipeptides, before the cyclization step, the first step is dehydration of Ser/Thr residues in the core peptide to generate dehydroalanine (Dha from Ser) or (Z)-dehydrobutyrine (Dhb from Thr). ^16^ Therefore, we next ran MD simulations starting from both unmodified peptides and dehydrated peptides to investigate whether the dehydration process affects the conformational preference of the peptides. Four possible initial rings can be formed in from the core peptide of ProcA3.3, connecting Thr11/Cys14, Thr11/Cys21, Cys14/Thr18 and Thr18/Cys21 residue pairs. Figure 3 shows the distance distribution of these four features for unmodified and fully dehydrated ProcA3.3 WT and Variant1. Unmodified and dehydrated peptide models demonstrate consistent distance distributions of all four features, indicating that the dehydration process does not greatly influence the predicted solution structure of the peptide in terms of favored conformations, although some changes in probability densities are observed.

**Figure 3:**
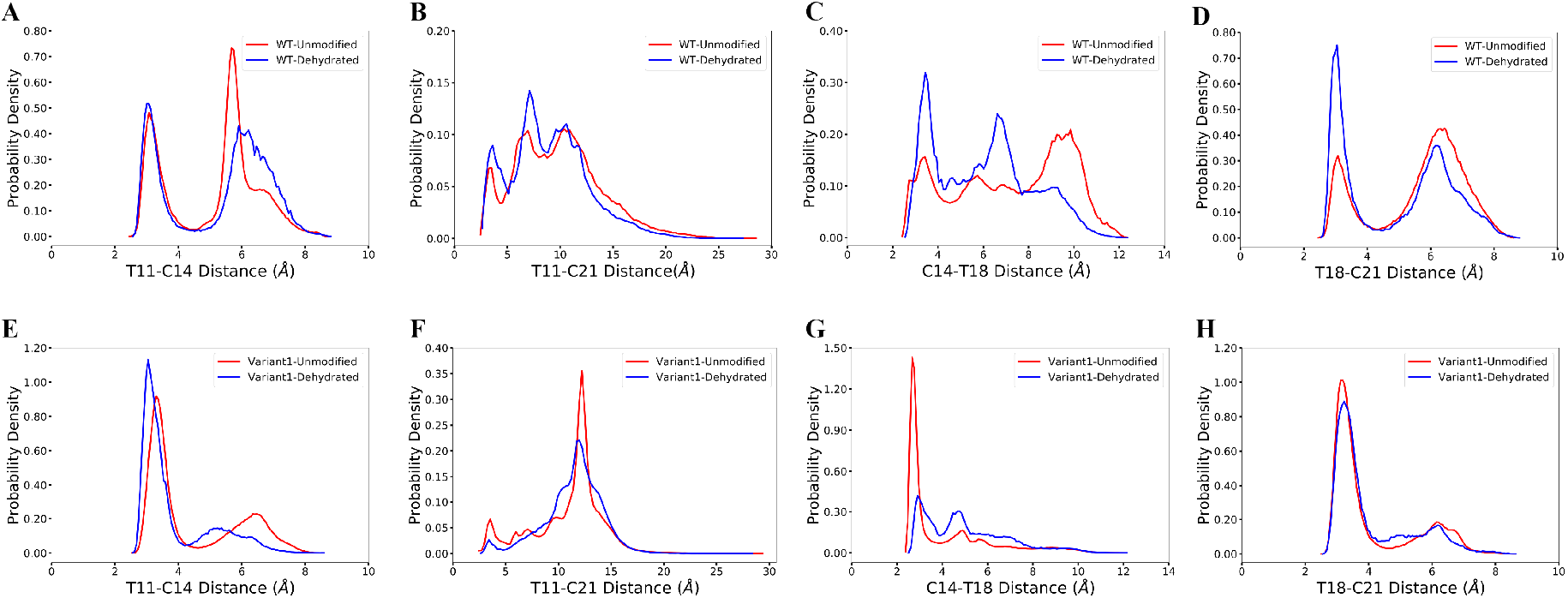
Comparison of unmodified and dehydrated ProcA3.3 variants. (A, B, C, D) Distance distribution of four important features for unmodified and fully dehydrated WT ProcA3.3. (E, F, G, H) Distance distribution of four important features for unmodified and fully dehydrated ProcA3.3 Variant1.

### Order of cyclization of prochlorosins

The cyclization process of prochlorosins is an attractive topic for investigation because for each substrate a single enzyme with one cyclization active site catalyzes the formation of a specific ring pattern out of many possible patterns. The formation of the final ring pattern is directly related to the order of cyclization since the first cyclization sets the final pattern. In the prochlorosin family, the three-dimensional structures of Pcn1.1, 2.1, 2.8, 2.10, 2.11 were determined by nuclear magnetic resonance (NMR) spectroscopy.^33^ Among them, both Pcn1.1 and 2.8 contain two non-overlapping rings, but the rings are of different sizes (Figure 4). Previous studies of ProcA2.8 have shown ring B (Ser13/Cys19) is formed first,^70^ where the correct formation of ring B is important for subsequent formation of ring A (Cys3/Ser9) via preorganization of the peptide structure to facilitate cyclization. ^74^ To explore whether substrate sequence can be utilized to predict the order of cyclization, we ran 25*μ*s MD simulation for ProcA2.8 in solution and calculated the probability of the formation of the four possible rings (Cys3/Ser9, Ser13/Cys19, Cys3/Ser13, Ser9/Cys19). Our result shows that ring B has a significantly higher probability to form (Figure 4A) and is more energetically favorable than ring A (Figure S5A). Thus, MD simulations mirror the experimental observation of ring B forming first.

**Figure 4:**
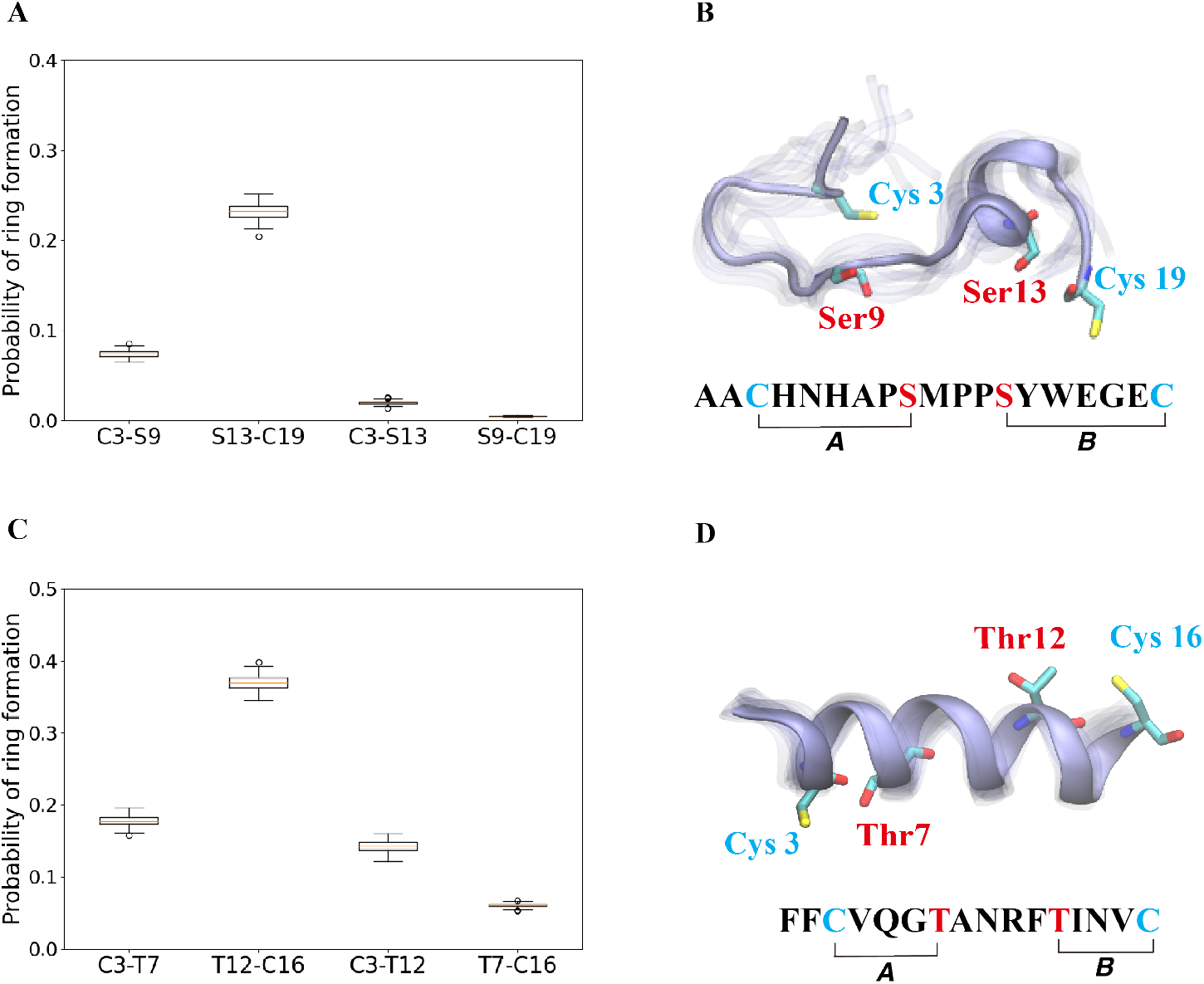
(A) Probability of formation of all possible rings for ProcA2.8, with ring formation defined as the distance between the *β*-carbon of Thr and the sulfur of Cys being within 7.5 Å. (B) Cartoon representation of the state leading to rings A and B in the free energy landscape for ProcA2.8 (for the entire free energy landscape of all states, see Figure S5A). In each state, 10 snapshots were randomly selected and aligned together. One snapshot is shown in ice blue, while the others are shown in transparent ice blue. (C) Probability of formation of all possible rings for ProcA1.1. (D) Cartoon representation of the state in which ring A and B are formed in the free energy landscape for ProcA1.1 (for the entire free energy landscape of all states, see Figure S5C). Error bar was calculated by 200 bootstrap samples with 80% of total trajectories.

Similarly, from the MSM weighted free energy landscape of ProcA1.1 (Figure S5C), we concluded that ring B (Thr12/Cys16) is more stable than ring A (Cys3/Thr7) and has a higher probability of formation (Figure 4C). To verify the order of cyclization predicted from simulation, ProcM was incubated with ProcA1.1 and the assay was quenched at various time points as described in the Methods section. After quenching, the samples were derivatized via NEM alkylation of free thiols to identify partially cyclized intermediates, and the leader peptide was removed using the protease LahT_150_ to obtain the core peptide. ^72^ Assay times and conditions were optimized to allow buildup of doubly dehydrated, singly cyclized intermediates. This peptide intermediate was then analyzed by LC-ESI MS/MS to obtain fragmentation data and determine the location(s) of NEM alkylation and thus cyclization.

The fragmentation data obtained of the monocyclized intermediate indicates Cys3 is alkylated and that ring B had been formed (Figure 5). These findings indicate that cyclization of Cys16 occurs before that of Cys3 and that the cyclization order of ProcA1.1 is similar to that of ProcA2.8.^69^ Notably, MD simulations correctly predicted the cyclization outcome. Importantly, C-to-N directionality is not uniform with ProcA substrates. For example, for WT ProcA3.3 the inner ring is formed first.^70^ Hence, the directionality is not enforced by the enzyme but is likely a result of the inherent reactivity of each Cys-Dha/Dhb pair, which in turn is governed by the conformational energy landscape.

**Figure 5:**
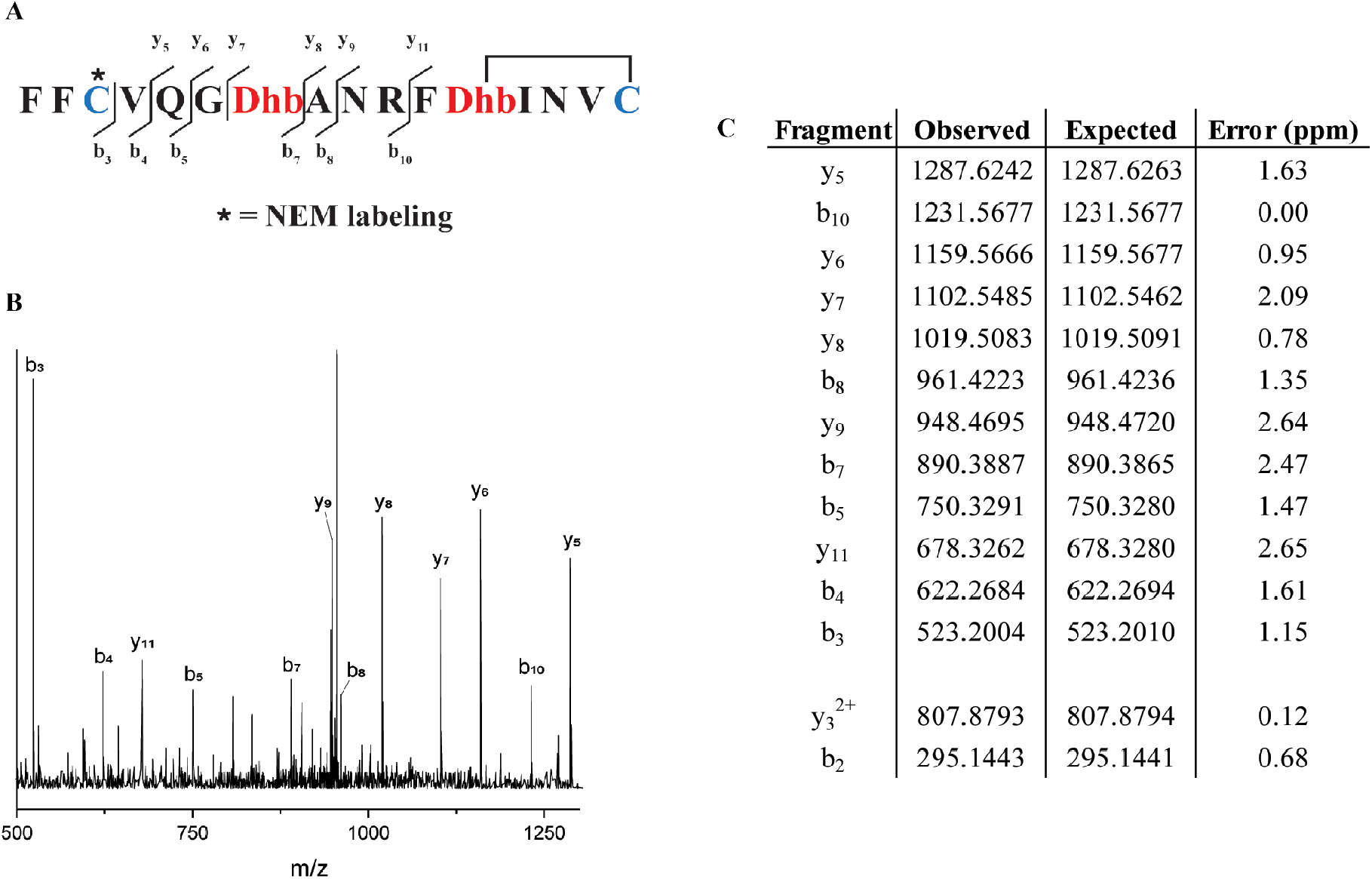
(A) Sequence of ProcA1.1 intermediate after leader peptide cleavage with observed fragmentation indicated. (B) Tandem-MS fragmentation of ProcA1.1. (C) Table of observed and expected fragment ion masses and the error calculated (ppm).

## Conclusion

In this work, we ran molecular dynamics simulation for the wild type substrate peptide ProcA3.3 and its variants in solution. Without the lanthipeptide synthetase, ProcM, as a catalyst, we predicted the same ring pattern for the core peptides of ProcA3.3 variants as the experimental results, with distance between reacting residues used as a proxy for the likelihood of a chemical reaction. Results from MD simulation provide an orthogonal line of evidence to support and explain how the core sequence can determine the final product’s ring pattern, rather than the enzyme. In addition, we demonstrate that in the biosynthetic pathway to prochlorosins, the dehydration step does not alter the predicted solution structure of the lanthipeptide. This finding suggests that even if the core sequence is not dehydrated, it can be used to determine the site-selectivity of the final products, an important finding for future prediction of ring patterns from sequence. Furthermore, we show that the secondary structure of the core peptide sequence is a crucial factor to control formation of the final product by determining the conformational energy landscape in solution.

Another interesting question we investigated in our work is the order of cyclization. We chose two candidates, ProcA1.1 and ProcA2.8 which both form two non-overlapping rings of different sizes, to see if we could utilize our simulations to predict the order of ring formation. Based on our MD simulation analysis of these two prochlorosins, the C-terminal ring has a higher probability to form and remain in a more energetically stable conformation, which suggests that the C-terminal ring will be first to form for both peptides. Previous study of ProcA2.8 indeed showed the C-terminal ring is the first ring to be formed during the biosynthesis by ProcM.^70^ In this study, we also experimentally verified the prediction from the MD simulation that ProcA1.1 is also cyclized in a C-to-N terminal order. Thus, for both ProcA1.1 and ProcA2.8, we see the same directionality of cyclization with and without ProcM as a catalyst, which provides further evidence to suggest substrate sequence controls the order of cyclization.

Overall, we utilized molecular dynamic simulation to demonstrate that the substrate core peptide sequence likely controls the final ring pattern of the prochlorosins and the order of the ring formation through determining secondary structure. This study therefore aids in the understanding of the mechanism of ring formation in the Proc system (and likely the SyncM system^31^) and the relationship between sequence and final product. In addition, our work may provide an effective strategy to facilitate the prediction of the ring pattern of lanthipeptides discovered by genome mining by analyzing the secondary structure of the core sequence and the conformational energy landscape. Compared to experimental investigations, the computational methodology is much faster and can accelerate bioengineering efforts and the design of novel lanthipeptides.

## Supporting information

Supplementary Images and Tables

## Acknowledgement

This work was supported by the National Institutes of Health grants R37 GM058822 to W.A.V. and R35 GM142745 to D.S. The authors thank Blue Water Supercomputing facility and Delta Supercomputer at National Center for Supercomputing Applications at University of Illinois Urbana-Champaign for providing the computational resources for this study.

## Supporting Information Available

Variation of Thr11-Cys14 distance projected along tIC1 and tIC2 for ProcA3.3 variants; MSM weighted free energy landscape projected along *α*−helix content located from Thr11 to Ala23 and Thr11-Cys14 distance for ProcA3.3 variants; Probability of formation of ring A′ (Cys14/Thr18) and B′ (Thr11/Cys21) for ProcA3.3 variants; Probability of formation of ring A′ (Thr11/Cys14) and B′ (Thr18/Cys21) for ProcA3.3 variants; MSM weighted free energy landscape for ProcA2.8 and ProcA1.1; Probability of formation of all possible rings for ProcA2.8 and ProcA1.1; Markov state model construction and analysis for ProcA3.3 WT; Markov state model construction and analysis for ProcA3.3 Variant1; Markov state model construction and analysis for ProcA3.3 Variant2; Markov state model construction and analysis for dehydrated ProcA3.3 WT; Markov state model construction and analysis for dehydrated ProcA3.3 Variant1; Markov state model construction and analysis for ProcA1.1; Markov state model construction and analysis for ProcA2.8; Chromatogram of ProcA1.1 purified by high-performance liquid chromatography; MALDI-TOF mass spectra of HPLC purified unmodified ProcA1.1; Table for Molecular Dynamics Simulation Systems.

## Data Availability

The data and code can be found at the following GitHub repository: https://github.com/ShuklaGroup/Lanthipeptide.

## References

(1) Newman, D. J.; Cragg, G. M. Natural Products as Sources of New Drugs over the Nearly Four Decades from 01/1981 to 09/2019. Journal of Natural Products 2020, 83, 770–803.

(2) Hetrick, K. J.; van der Donk, W. A. Ribosomally synthesized and post-translationally modified peptide natural product discovery in the genomic era. Current Opinion in Chemical Biology 2017, 38, 36–44.

(3) Luo, S.; Dong, S.-H. Recent advances in the discovery and biosynthetic study of eukaryotic RiPP natural products. Molecules 2019, 24, 1541.

(4) Scherlach, K.; Hertweck, C. Mining and unearthing hidden biosynthetic potential. Nature Communications 2021, 12, 3864.

(5) Medema, M. H.; de Rond, T.; Moore, B. S. Mining genomes to illuminate the specialized chemistry of life. Nature Reviews Genetics 2021, 22, 553–571.

(6) Albarano, L.; Esposito, R.; Ruocco, N.; Costantini, M. Genome mining as new challenge in natural products discovery. Marine drugs 2020, 18, 199.

(7) Baltz, R. H. Genome mining for drug discovery: Progress at the front end. Journal of Industrial Microbiology and Biotechnology 2021, 48, kuab044.

(8) Kenshole, E.; Herisse, M.; Michael, M.; Pidot, S. J. Natural product discovery through microbial genome mining. Current Opinion in Chemical Biology 2021, 60, 47–54.

(9) Arnison, P. G.; Bibb, M. J.; Bierbaum, G.; Bowers, A. A.; Bugni, T. S.; Bulaj, G.; Camarero, J. A.; Campopiano, D. J.; Challis, G. L.; Clardy, J.; Cotter, P. D.; Craik, D. J.; Dawson, M.; Dittmann, E.; Donadio, S.; Dorrestein, P. C.; Entian, K.-D.; Fischbach, M. A.; Garavelli, J. S.; Göransson, U.; Gruber, C. W.; Haft, D. H.; Hemscheidt, T. K.; Hertweck, C.; Hill, C.; Horswill, A. R.; Jaspars, M.; Kelly, W. L.; Klinman, J. P.; Kuipers, O. P.; Link, A. J.; Liu, W.; Marahiel, M. A.; Mitchell, D. A.; Moll, G. N.; Moore, B. S.; Müller, R.; Nair, S. K.; Nes, I. F.; Norris, G. E.; Olivera, B. M.; Onaka, H.; Patchett, M. L.; Piel, J.; Reaney, M. J. T.; Rebuffat, S.; Ross, R. P.; Sahl, H.-G.; Schmidt, E. W.; Selsted, M. E.; Severinov, K.; Shen, B.; Sivonen, K.; Smith, L.; Stein, T.; Süssmuth, R. D.; Tagg, J. R.; Tang, G.-L.; Truman, A. W.; Vederas, J. C.; Walsh, C. T.; Walton, J. D.; Wenzel, S. C.; Willey, J. M.; van der Donk, W. A. Ribosomally synthesized and post-translationally modified peptide natural products: overview and recommendations for a universal nomenclature. Natural product reports 2013, 30, 108–160.

(10) Walker, M. C.; Eslami, S. M.; Hetrick, K. J.; Ackenhusen, S. E.; Mitchell, D. A.; van der Donk, W. A. Precursor peptide-targeted mining of more than one hundred thousand genomes expands the lanthipeptide natural product family. BMC Genomics 2020, 21, 387.

(11) Montalbán-López, M.; Scott, T. A.; Ramesh, S.; Rahman, I. R.; van Heel, A. J.; Viel, J. H.; Bandarian, V.; Dittmann, E.; Genilloud, O.; Goto, Y.; Burgos, M. J. G.; Hill, C.; Kim, S.; Koehnke, J.; Latham, J. A.; Link, A. J.; Martínez, B.; Nair, S. K.; Nicolet, Y.; Rebuffat, S.; Sahl, H.-G.; Sareen, D.; Schmidt, E. W.; Schmitt, L.; Severinov, K.; Süssmuth, R. D.; Truman, A. W.; Wang, H.; Weng, J.-K.; van Wezel, G. P.; Zhang, Q.; Zhong, J.; Piel, J.; Mitchell, D. A.; Kuipers, O. P.; van der Donk, W. A. New developments in RiPP discovery, enzymology and engineering. Natural Product Reports 2021, 38, 130–239.

(12) Lubelski, J.; Rink, R.; Khusainov, R.; Moll, G. N.; Kuipers, O. P. Biosynthesis, immunity, regulation, mode of action and engineering of the model lantibiotic nisin. Cellular and Molecular Life Sciences 2007, 65, 455–476.

(13) Smith, T. E.; Pond, C. D.; Pierce, E.; Harmer, Z. P.; Kwan, J.; Zachariah, M. M.; Harper, M. K.; Wyche, T. P.; Matainaho, T. K.; Bugni, T. S.; Barrows, L. R.; Ireland, C. M.; Schmidt, E. W. Accessing chemical diversity from the uncultivated symbionts of small marine animals. Nature Chemical Biology 2018, 14, 179–185.

(14) Mohr, K. I.; Volz, C.; Jansen, R.; Wray, V.; Hoffmann, J.; Bernecker, S.; Wink, J.; Gerth, K.; Stadler, M.; Müller, R. Pinensins: The First Antifungal Lantibiotics. Angewandte Chemie International Edition 2015, 54, 11254–11258.

(15) Meindl, K.; Schmiederer, T.; Schneider, K.; Reicke, A.; Butz, D.; Keller, S.; Gühring, H.; Vértesy, L.; Wink, J.; Hoffmann, H.; Brönstrup, M.; Sheldrick, G.; Süssmuth, R. Labyrinthopeptins: A New Class of Carbacyclic Lantibiotics. Angewandte Chemie International Edition 2010, 49, 1151–1154.

(16) Repka, L. M.; Chekan, J. R.; Nair, S. K.; van der Donk, W. A. Mechanistic Understanding of Lanthipeptide Biosynthetic Enzymes. Chemical Reviews 2017, 117, 5457–5520.

(17) Field, D.; Collins, B.; Cotter, P. D.; Hill, C.; Ross, R. P. A System for the Random Mutagenesis of the Two-Peptide Lantibiotic Lacticin 3147: Analysis of Mutants Producing Reduced Antibacterial Activities. Journal of Molecular Microbiology and Biotechnology 2007, 13, 226–234.

(18) Caetano, T.; Krawczyk, J. M.; Mösker, E.; Süssmuth, R. D.; Mendo, S. Heterologous Expression, Biosynthesis, and Mutagenesis of Type II Lantibiotics from Bacillus licheniformis in Escherichia coli. Chemistry & Biology 2011, 18, 90–100.

(19) Chen, S.; Wilson-Stanford, S.; Cromwell, W.; Hillman, J. D.; Guerrero, A.; Allen, C. A.; Sorg, J. A.; Smith, L. Site-Directed Mutations in the Lanthipeptide Mutacin 1140. Applied and Environmental Microbiology 2013, 79, 4015–4023.

(20) Vinogradov, A. A.; Chang, J. S.; Onaka, H.; Goto, Y.; Suga, H. Accurate Models of Substrate Preferences of Post-Translational Modification Enzymes from a Combination of mRNA Display and Deep Learning. ACS Central Science 2022, 8, 814–824.

(21) Field, D.; Cotter, P. D.; Hill, C.; Ross, R. P. Bioengineering lantibiotics for therapeutic success. Frontiers in microbiology 2015, 6, 1363.

(22) Montalbán-López, M.; van Heel, A. J.; Kuipers, O. P. Employing the promiscuity of lantibiotic biosynthetic machineries to produce novel antimicrobials. FEMS Microbiology Reviews 2016, 41, 5–18.

(23) Kuthning, A.; Durkin, P.; Oehm, S.; Hoesl, M. G.; Budisa, N.; Süssmuth, R. D. Towards Biocontained Cell Factories: An Evolutionarily Adapted Escherichia coliStrain Produces a New-to-nature Bioactive Lantibiotic ContainingThienopyrrole-Alanine. Scientific reports 2016, 6, 33447.

(24) Burkhart, B. J.; Kakkar, N.; Hudson, G. A.; van der Donk, W. A.; Mitchell, D. A. Chimeric Leader Peptides for the Generation of Non-Natural Hybrid RiPP Products. ACS Central Science 2017, 3, 629–638.

(25) Hetrick, K. J.; Walker, M. C.; van der Donk, W. A. Development and Application of Yeast and Phage Display of Diverse Lanthipeptides. ACS Central Science 2018, 4, 458–467.

(26) Yang, X.; Lennard, K. R.; He, C.; Walker, M. C.; Ball, A. T.; Doigneaux, C.; Tavassoli, A.; van der Donk, W. A. A lanthipeptide library used to identify a protein–protein interaction inhibitor. Nature Chemical Biology 2018, 14, 375–380.

(27) Schmitt, S.; Montalbán-López, M.; Peterhoff, D.; Deng, J.; Wagner, R.; Held, M.; Kuipers, O. P.; Panke, S. Analysis of modular bioengineered antimicrobial lanthipeptides at nanoliter scale. Nature Chemical Biology 2019, 15, 437–443.

(28) Moll, G. N.; Kuipers, A.; Rink, R.; Bosma, T.; de Vries, L.; Namsolleck, P. Biosynthesis of lanthionine-constrained agonists of G protein-coupled receptors. Biochemical Society Transactions 2020, 48, 2195–2203.

(29) Li, B.; Sher, D.; Kelly, L.; Shi, Y.; Huang, K.; Knerr, P. J.; Joewono, I.; Rusch, D.; Chisholm, S. W.; van der Donk, W. A. Catalytic promiscuity in the biosynthesis of cyclic peptide secondary metabolites in planktonic marine cyanobacteria. Proceedings of the National Academy of Sciences 2010, 107, 10430–10435.

(30) Cubillos-Ruiz, A.; Berta-Thompson, J. W.; Becker, J. W.; van der Donk, W. A.; Chisholm, S. W. Evolutionary radiation of lanthipeptides in marine cyanobacteria. Proceedings of the National Academy of Sciences 2017, 114, E5424–E5433.

(31) Arias-Orozco, P.; Inklaar, M.; Lanooij, J.; Cebrián, R.; Kuipers, O. P. Functional Expression and Characterization of the Highly Promiscuous Lanthipeptide Synthetase SyncM, Enabling the Production of Lanthipeptides with a Broad Range of Ring Topologies. ACS Synthetic Biology 2021, 10, 2579–2591.

(32) Tang, W.; van der Donk, W. A. Structural Characterization of Four Prochlorosins: A Novel Class of Lantipeptides Produced by Planktonic Marine Cyanobacteria. Biochemistry 2012, 51, 4271–4279.

(33) Bobeica, S. C.; Zhu, L.; Acedo, J. Z.; Tang, W.; van der Donk, W. A. Structural determinants of macrocyclization in substrate-controlled lanthipeptide biosynthetic pathways. Chemical Science 2020, 11, 12854–12870.

(34) Le, T.; Fouque, K. J. D.; Santos-Fernandez, M.; Navo, C. D.; Jiménez-Osés, G.; Sarksian, R.; Fernandez-Lima, F. A.; van der Donk, W. A. Substrate Sequence Controls Regioselectivity of Lanthionine Formation by ProcM. Journal of the American Chemical Society 2021, 143, 18733–18743.

(35) Yu, Y.; Zhang, Q.; van der Donk, W. A. Insights into the evolution of lanthipeptide biosynthesis. Protein Science 2013, 22, 1478–1489.

(36) Le, T.; van der Donk, W. A. Mechanisms and Evolution of Diversity-Generating RiPP Biosynthesis. Trends in Chemistry 2021, 3, 266–278.

(37) Lindorff-Larsen, K.; Piana, S.; Dror, R. O.; Shaw, D. E. How fast-folding proteins fold. Science 2011, 334, 517–520.

(38) Piana, S.; Klepeis, J. L.; Shaw, D. E. Assessing the accuracy of physical models used in protein-folding simulations: quantitative evidence from long molecular dynamics simulations. Current opinion in structural biology 2014, 24, 98–105.

(39) Kuroda, Y.; Suenaga, A.; Sato, Y.; Kosuda, S.; Taiji, M. All-atom molecular dynamics analysis of multi-peptide systems reproduces peptide solubility in line with experimental observations. Scientific reports 2016, 6, 19479.

(40) Shrestha, U. R.; Juneja, P.; Zhang, Q.; Gurumoorthy, V.; Borreguero, J. M.; Urban, V.; Cheng, X.; Pingali, S. V.; Smith, J. C.; O’Neill, H. M., et al. Generation of the configurational ensemble of an intrinsically disordered protein from unbiased molecular dynamics simulation. Proceedings of the National Academy of Sciences 2019, 116, 20446–20452.

(41) Damjanovic, J.; Miao, J.; Huang, H.; Lin, Y.-S. Elucidating solution structures of cyclic peptides using molecular dynamics simulations. Chemical Reviews 2021, 121, 2292–2324.

(42) Schrödinger, L.; DeLano, W. PyMOL. http://www.pymol.org/pymol.

(43) Martínez, L.; Andrade, R.; Birgin, E.; Martínez, J. M. PACKMOL: A package for building initial configurations for molecular dynamics simulations. Journal of Computational Chemistry 2009, 30, 2157 – 2164.

(44) Case, D.; Belfon, K.; Ben-Shalom, I.; Brozell, S.; Cerutti, D.; Cheatham III, T.; Cruzeiro, V.; Darden, T.; R.E., D. AMBER 2018.

(45) Bayly, C. I.; Cieplak, P.; Cornell, W.; Kollman, P. A. A well-behaved electrostatic potential based method using charge restraints for deriving atomic charges: the RESP model. The Journal of Physical Chemistry 1993, 97, 10269–10280.

(46) Frisch, M. J.; Trucks, G. W.; Schlegel, H. B.; Scuseria, G. E.; Robb, M. A.; Cheeseman, J. R.; Scalmani, G.; Barone, V.; Petersson, G. A.; Nakatsuji, H.; Li, X.; Caricato, M.; Marenich, A. V.; Bloino, J.; Janesko, B. G.; Gomperts, R.; Mennucci, B.; Hratchian, H. P.; Ortiz, J. V.; Izmaylov, A. F.; Sonnenberg, J. L.; Williams-Young, D.; Ding, F.; Lipparini, F.; Egidi, F.; Goings, J.; Peng, B.; Petrone, A.; Henderson, T.; Ranasinghe, D.; Zakrzewski, V. G.; Gao, J.; Rega, N.; Zheng, G.; Liang, W.; Hada, M.; Ehara, M.; Toyota, K.; Fukuda, R.; Hasegawa, J.; Ishida, M.; Nakajima, T.; Honda, Y.; Kitao, O.; Nakai, H.; Vreven, T.; Throssell, K.; Montgomery, J. A., Jr.; Peralta, J. E.; Ogliaro, F.; Bearpark, M. J.; Heyd, J. J.; Brothers, E. N.; Kudin, K. N.; Staroverov, V. N.; Keith, T. A.; Kobayashi, R.; Normand, J.; Raghavachari, K.; Rendell, A. P.; Burant, J. C.; Iyengar, S. S.; Tomasi, J.; Cossi, M.; Millam, J. M.; Klene, M.; Adamo, C.; Cammi, R.; Ochterski, J. W.; Martin, R. L.; Morokuma, K.; Farkas, O.; Foresman, J. B.; Fox, D. J. Gaussian 16 Revision C.01. 2016; Gaussian Inc. Wallingford CT.

(47) Loncharich, R. J.; Brooks, B. R.; Pastor, R. W. Langevin dynamics of peptides: The frictional dependence of isomerization rates of N-acetylalanyl-N’-methylamide. Biopolymers 1992, 32, 523–535.

(48) Aqvist, J.; Wennerström, P.; Nervall, M.; Bjelic, S.; Brandsdal, B. Molecular dynamics simulations of water and biomolecules wit a Monte Carlo constant pressure algorithm. Chemical Physics Letters 2004, 384, 288–294.

(49) Braun, E.; Gilmer, J.; Mayes, H. B.; Mobley, D. L.; Monroe, J. I.; Prasad, S.; Zuckerman, D. M. Best Practices for Foundations in Molecular Simulations [Article v1.0]. Living Journal of Computational Molecular Science 2019, 1, 5957.

(50) Ryckaert, J.-P.; Ciccotti, G.; Berendsen, H. J. Numerical integration of the cartesian equations of motion of a system with constraints: molecular dynamics of n-alkanes. Journal of Computational Physics 1977, 23, 327–341.

(51) Hopkins, C. W.; Grand, S. L.; Walker, R. C.; Roitberg, A. E. Long-Time-Step Molecular Dynamics through Hydrogen Mass Repartitioning. Journal of Chemical Theory and Computation 2015, 11, 1864–1874.

(52) Bowman, G. R.; Ensign, D. L.; Pande, V. S. Enhanced Modeling via Network Theory: Adaptive Sampling of Markov State Models. Journal of Chemical Theory and Computation 2010, 6, 787–794.

(53) Lawrenz, M.; Shukla, D.; Pande, V. S. Cloud computing approaches for prediction of ligand binding poses and pathways. Scientific reports 2015, 5, 7918.

(54) Lane, T. J.; Shukla, D.; Beauchamp, K. A.; Pande, V. S. To milliseconds and beyond: challenges in the simulation of protein folding. Current Opinion in Structural Biology 2013, 23, 58–65.

(55) Dutta, S.; Selvam, B.; Shukla, D. Distinct Binding Mechanisms for Allosteric Sodium Ion in Cannabinoid Receptors. ACS Chemical Neuroscience 2022, 13, 379–389.

(56) Dutta, S.; Selvam, B.; Das, A.; Shukla, D. Mechanistic origin of partial agonism of tetrahydrocannabinol for cannabinoid receptors. Journal of Biological Chemistry 2022, 298, 101764.

(57) Sculley, D. Web-scale k-means clustering.Proceedings of the 19th international conference on World wide web - WWW ‘10. 2010.

(58) Shukla, D.; Hernández, C. X.; Weber, J. K.; Pande, V. S. Markov State Models Provide Insights into Dynamic Modulation of Protein Function. Accounts of Chemical Research 2015, 48, 414–422.

(59) Husic, B. E.; Pande, V. S. Markov State Models: From an Art to a Science. Journal of the American Chemical Society 2018, 140, 2386–2396.

(60) Sidky, H.; Chen, W.; Ferguson, A. L. High-Resolution Markov State Models for the Dynamics of Trp-Cage Miniprotein Constructed Over Slow Folding Modes Identified by State-Free Reversible VAMPnets. The Journal of Physical Chemistry B 2019, 123, 7999–8009.

(61) Wu, H.; Noé, F. Variational Approach for Learning Markov Processes from Time Series Data. Journal of Nonlinear Science 2019, 30, 23–66.

(62) Pérez-Hernández, G.; Paul, F.; Giorgino, T.; Fabritiis, G. D.; Noé, F. Identification of slow molecular order parameters for Markov model construction. The Journal of Chemical Physics 2013, 139, 015102.

(63) Noé, F.; Clementi, C. Kinetic Distance and Kinetic Maps from Molecular Dynamics Simulation. Journal of Chemical Theory and Computation 2015, 11, 5002–5011.

(64) Scherer, M. K.; Trendelkamp-Schroer, B.; Paul, F.; Pérez-Hernández, G.; Hoffmann, M.; Plattner, N.; Wehmeyer, C.; Prinz, J.-H.; Noé, F. PyEMMA 2: A Software Package for Estimation, Validation, and Analysis of Markov Models. Journal of Chemical Theory and Computation 2015, 11, 5525–5542.

(65) McGibbon, R. T.; Beauchamp, K. A.; Harrigan, M. P.; Klein, C.; Swails, J. M.; Hernández, C. X.; Schwantes, C. R.; Wang, L.-P.; Lane, T. J.; Pande, V. S. MD-Traj: A Modern Open Library for the Analysis of Molecular Dynamics Trajectories. Biophysical Journal 2015, 109, 1528–1532.

(66) Humphrey, W.; Dalke, A.; Schulten, K. VMD: Visual molecular dynamics. Journal of Molecular Graphics 1996, 14, 33–38.

(67) Hunter, J. D. Matplotlib: A 2D Graphics Environment. Computing in Science & Engineering 2007, 9, 90–95.

(68) Sultan, M. M.; Kiss, G.; Pande, V. S. Towards simple kinetic models of functional dynamics for a kinase subfamily. Nature Chemistry 2018, 10, 903–909.

(69) Thibodeaux, C. J.; Ha, T.; van der Donk, W. A. A Price To Pay for Relaxed Substrate Specificity: A Comparative Kinetic Analysis of the Class II Lanthipeptide Synthetases ProcM and HalM2. Journal of the American Chemical Society 2014, 136, 17513–17529.

(70) Mukherjee, S.; van der Donk, W. A. Mechanistic Studies on the Substrate-Tolerant Lanthipeptide Synthetase ProcM. Journal of the American Chemical Society 2014, 136, 10450–10459.

(71) Bobeica, S. C.; van der Donk, W. A. Methods in Enzymology; Elsevier, 2018; pp 165– 203.

(72) Bobeica, S. C.; Dong, S.-H.; Huo, L.; Mazo, N.; McLaughlin, M. I.; Jiménez-Osés, G.; Nair, S. K.; van der Donk, W. A. Insights into AMS/PCAT transporters from biochemical and structural characterization of a double Glycine motif protease. eLife 2019, 8, e42305.

(73) Yu, Y.; Mukherjee, S.; van der Donk, W. A. Product Formation by the Promiscuous Lanthipeptide Synthetase ProcM is under Kinetic Control. Journal of the American Chemical Society 2015, 137, 5140–5148.

(74) Hegemann, J. D.; Bobeica, S. C.; Walker, M. C.; Bothwell, I. R.; van der Donk, W. A. Assessing the Flexibility of the Prochlorosin 2.8 Scaffold for Bioengineering Applications. ACS Synthetic Biology 2019, 8, 1204–1214.

