## Supplementary Images and Tables for "Sequence Controlled Secondary Structure Determines Site-selectivity of Lanthipeptides"

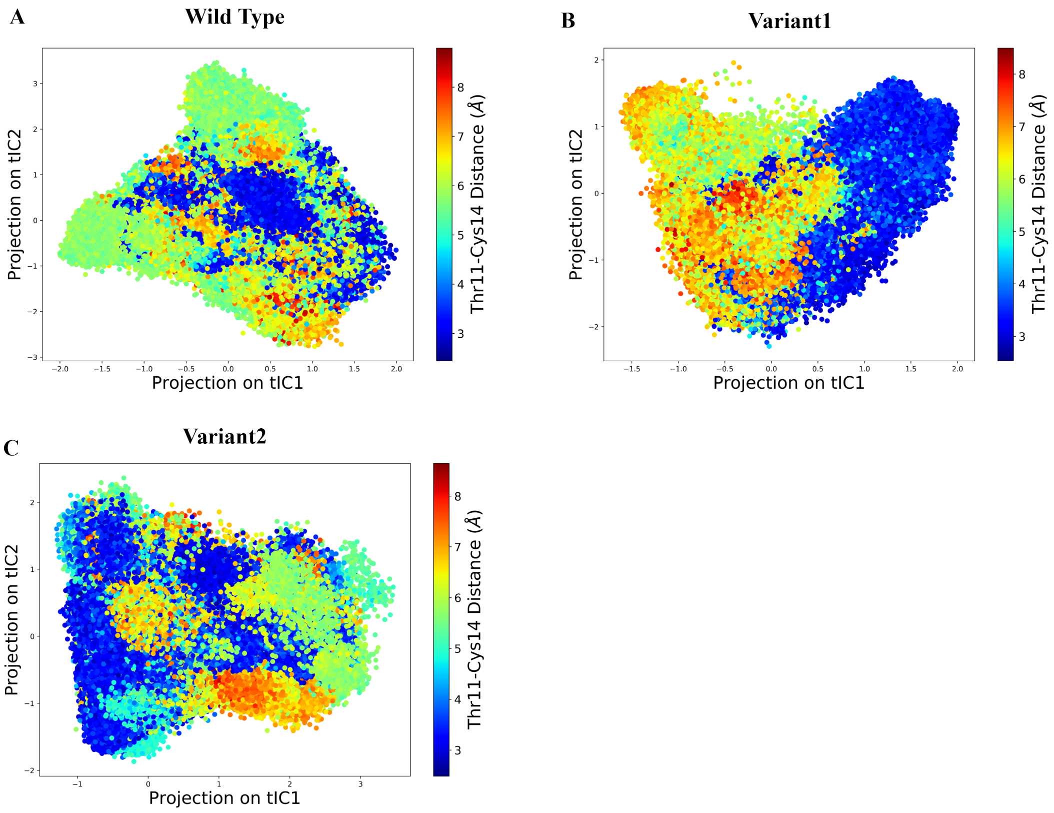


Figure S1: (A, B, C) Variation of Thr11-Cys14 distance projected along tIC1 and tIC2 for ProcA3.3 core peptide variants. tIC1 and tIC2 are the first and second component of time-lagged independent component analysis (described in the Method section). They represent the two slowest components in the system. Thr11-Cys14 distance has a high correlation with tIC1, especially for the ProcA3.3 Variant1.

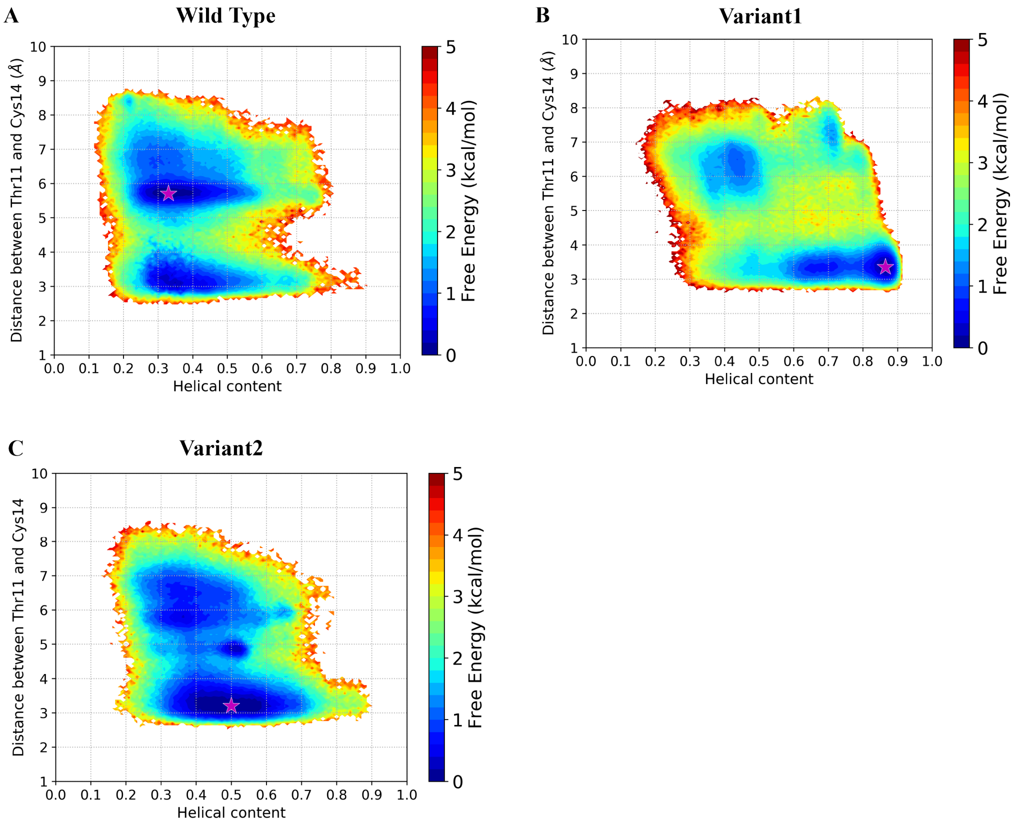


Figure S2: (A, B, C) MSM (Markov state model) weighted free energy landscape projected along $\alpha$-helix content (located from Thr11 to Ala23) and Thr11-Cys14 distance for ProcA3.3 variants. Star region is the most populated state in the system.

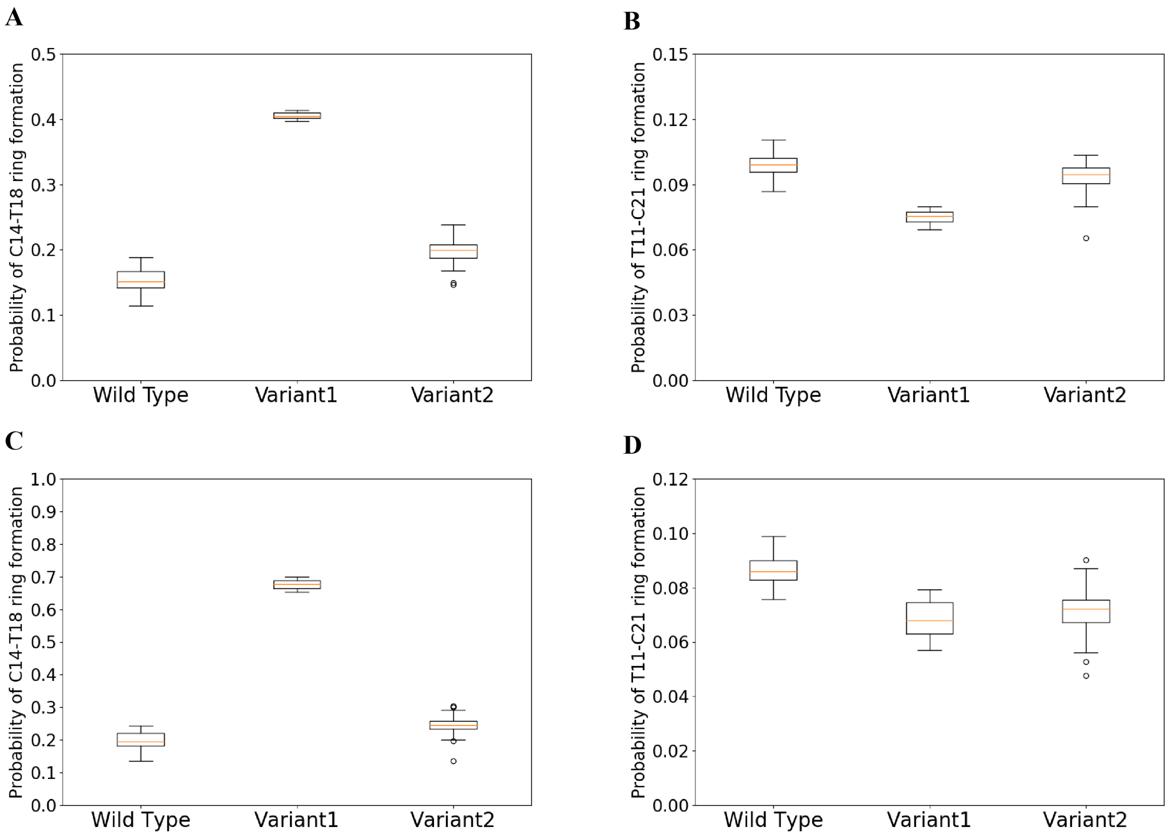


Figure S3: (A, B) Probability of formation of ring A′ (Cys14/Thr18) and B′ (Thr11/Cys21) for ProcA3.3 variants, with ring formation defined as the distance between the $\beta$-carbon of Thr and the sulfur of Cys being within 7.5 Å. (C, D) Probability of formation of ring A′ (Cys14/Thr18) and B′ (Thr11/Cys21) for ProcA3.3 variants, with ring formation defined as the distance between any of their heavy atoms being within 4.5 Å. Error bar was calculated by 100 bootstrap samples with 80% of total trajectories.

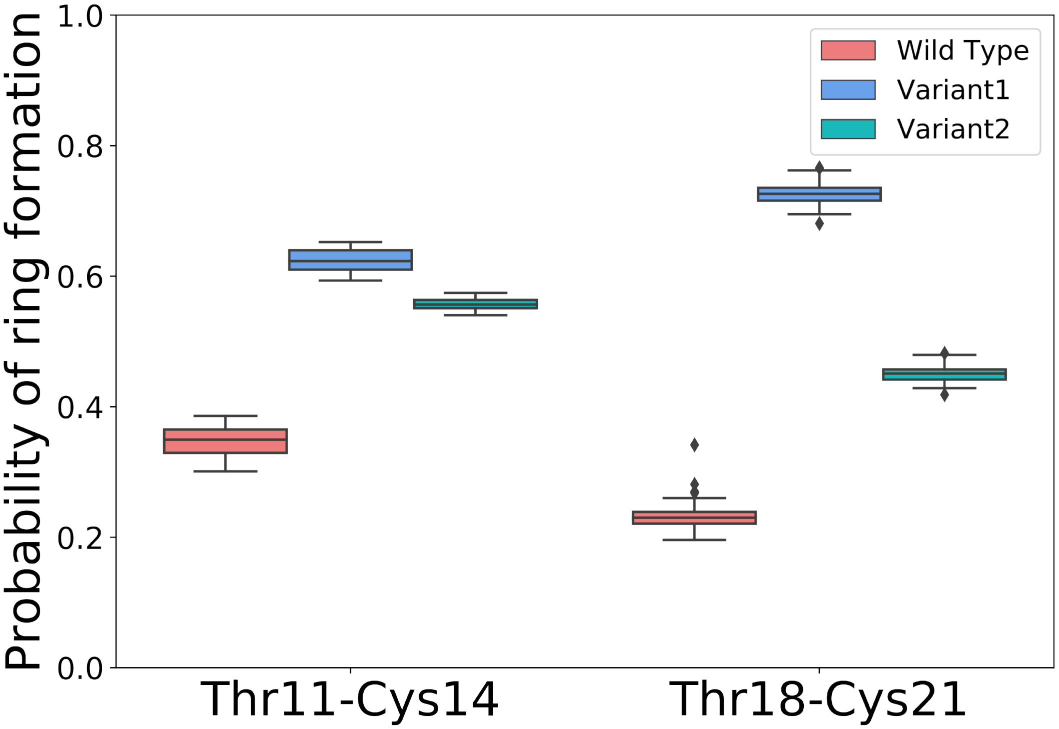


Figure S4: Probability of formation of ring A$'$ (Thr11/Cys14) and B$'$(Thr18/Cys21) for ProcA3.3 variants, with ring formation defined as the distance between any of their heavy atoms being within 4.5 Å. Error bar was calculated by 100 bootstrap samples with 80% of total trajectories.

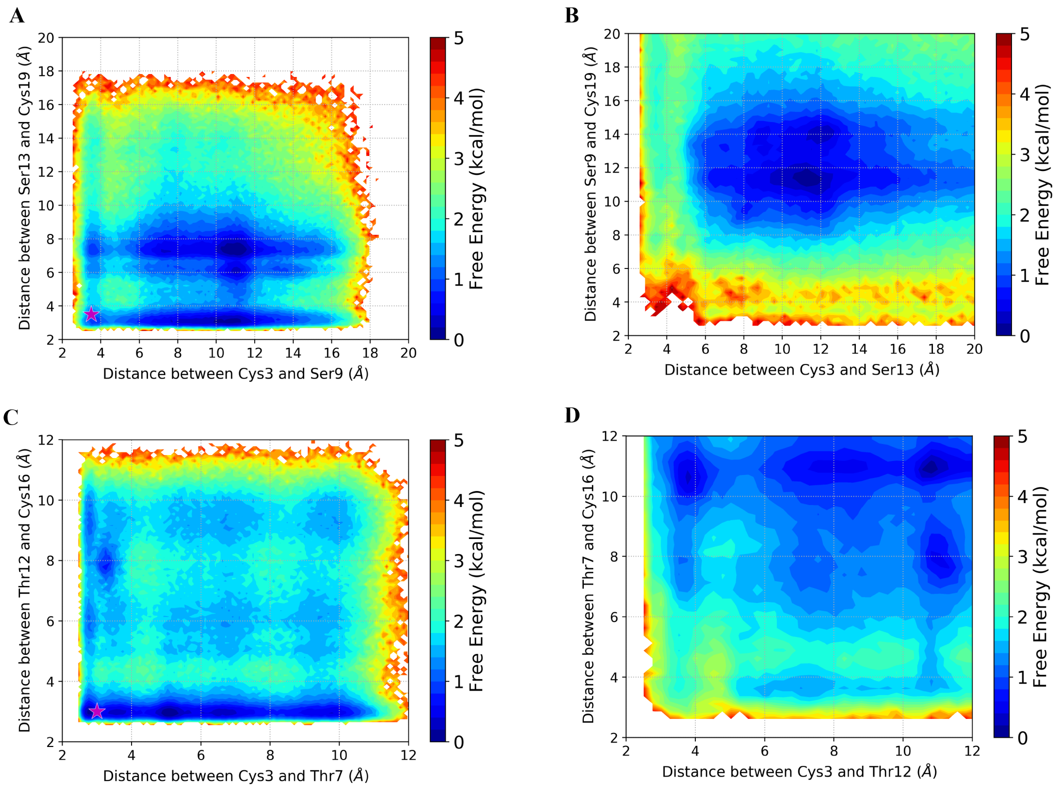


Figure S5: (A) MSM (Markov state model) weighted free energy landscape projected along the Cys3-Ser9 distance and the Ser13-Cys19 distance for ProcA2.8. (B) MSM weighted free energy landscape projected along the Cys3-Ser13 distance and the Ser9-Cys19 distance for ProcA2.8. (C) MSM weighted free energy landscape projected along the Cys3-Thr7 distance and the Thr12-Cys16 distance for ProcA1.1. (D) MSM weighted free energy landscape projected along the Cys3-Thr12 distance and the Thr7-Cys16 distance for ProcA1.1. Star region shows the state leading to rings A and B in the free energy landscape.

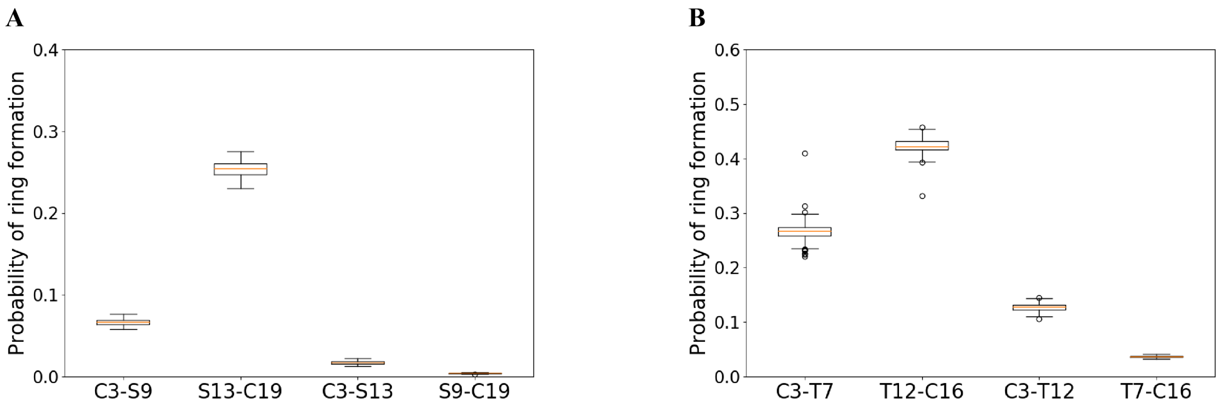


Figure S6: (A, B) Probability of formation of all possible rings for ProcA2.8 and ProcA1.1, with ring formation defined as the distance between any of their heavy atoms being within 4.5 Å. Error bar was calculated by 100 bootstrap samples with 80% of total trajectories.

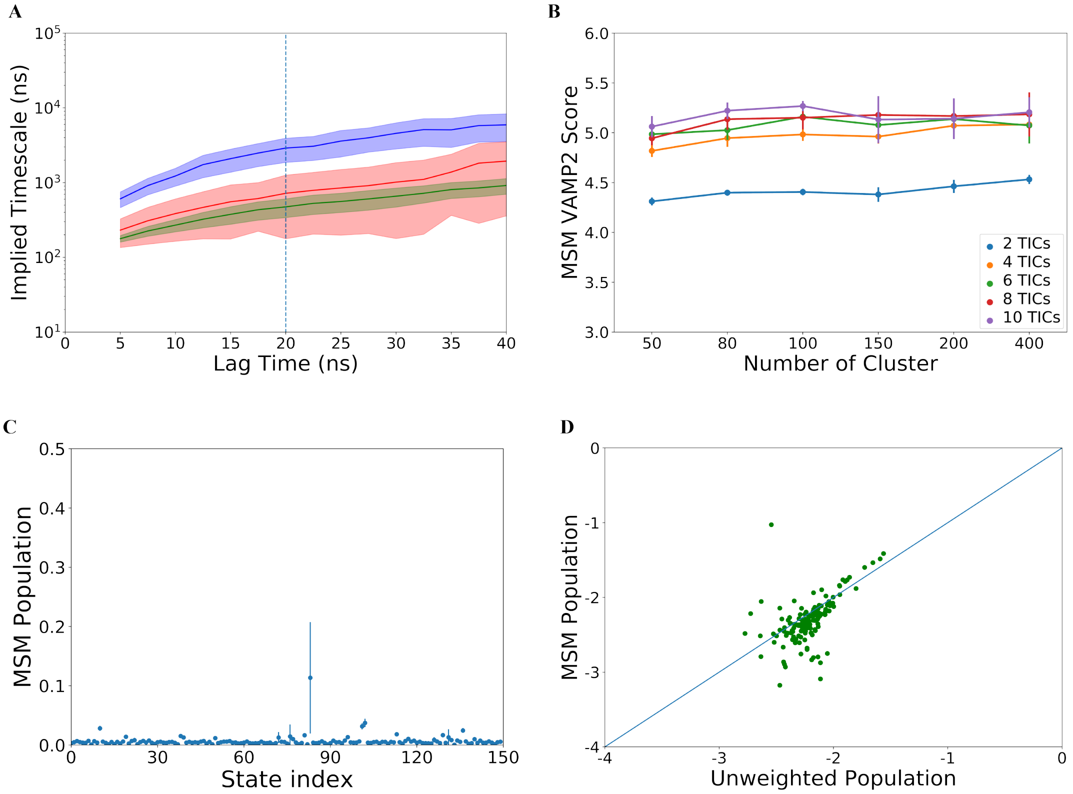


Figure S7: **Markov state model construction and analysis for ProcA3.3 WT.** (A) Implied timescale calculated from the 2nd (blue), 3rd (red) and 4th (green) highest MSM eigenvalues at different lag times, MSM lagtime of 20 ns was chosen. (B) VAMP-2 scores with different clusters and different number of tICs of MSM, 150 clusters and 10 tic dimensions were selected for MSM. (C) MSM population density of different MSM state index, error bar was calculated by 200 rounds bootstrapping with 80% of total trajectories. (D) MSM population vs Raw population in each state.

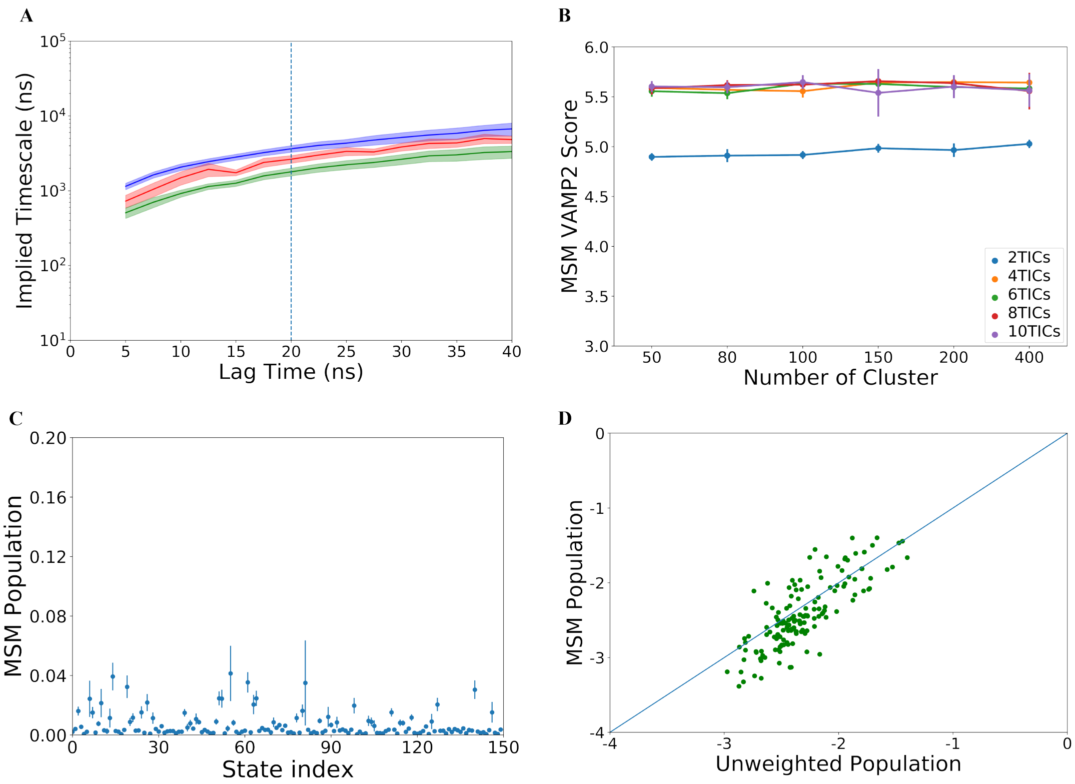


Figure S8: **Markov state model construction and analysis for ProcA3.3 Variant1.** (A) Implied timescale calculated from the 2nd (blue), 3rd (red) and 4th (green) highest MSM eigenvalues at different lag times, MSM lagtime of 20 ns was chosen. (B) VAMP-2 scores with different clusters and different number of tICs of MSM, 150 clusters and 8 tic dimensions were selected for MSM. (C) MSM population density of different MSM state index, error bar was calculated by 200 rounds bootstrapping with 80% of total trajectories. (D) MSM population vs Raw population in each state.

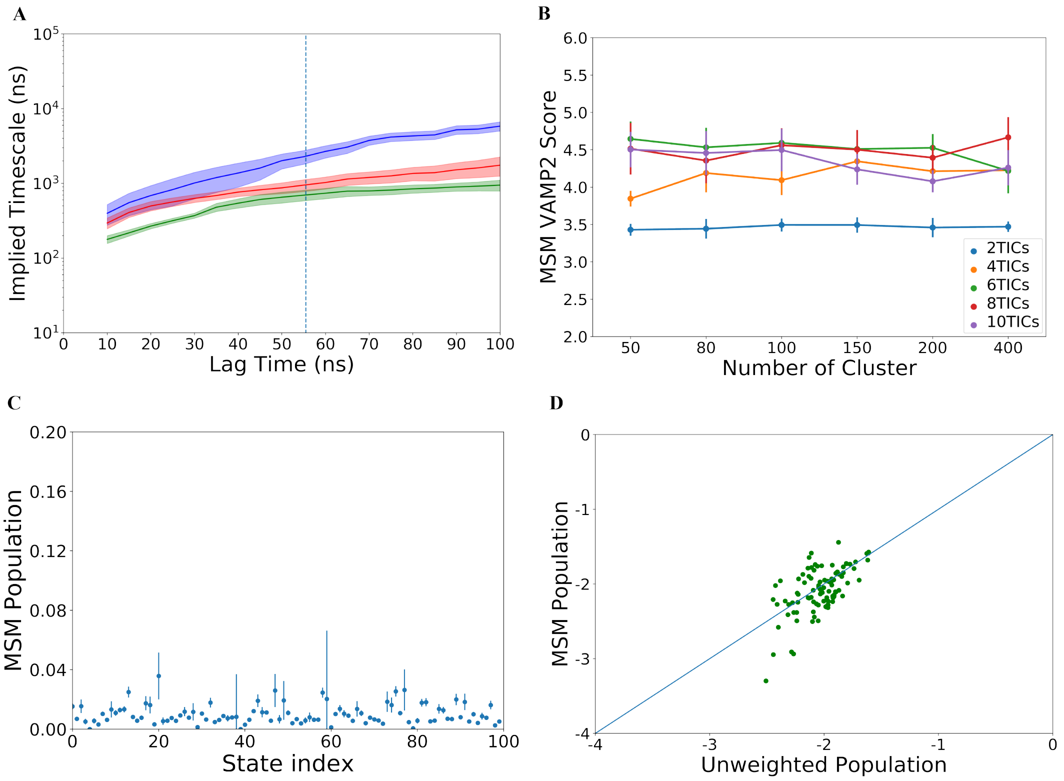


Figure S9: **Markov state model construction and analysis for ProcA3.3 Variant2.** (A) Implied timescale calculated from the 2nd (blue), 3rd (red) and 4th (green) highest MSM eigenvalues at different lag times, MSM lagtime of 55 ns was chosen. (B) VAMP-2 scores with different clusters and different number of tICs of MSM, 100 clusters and 6 tic dimensions were selected for MSM. (C) MSM population density of different MSM state index, error bar was calculated by 200 rounds bootstrapping with 80% of total trajectories. (D) MSM population vs Raw population in each state.

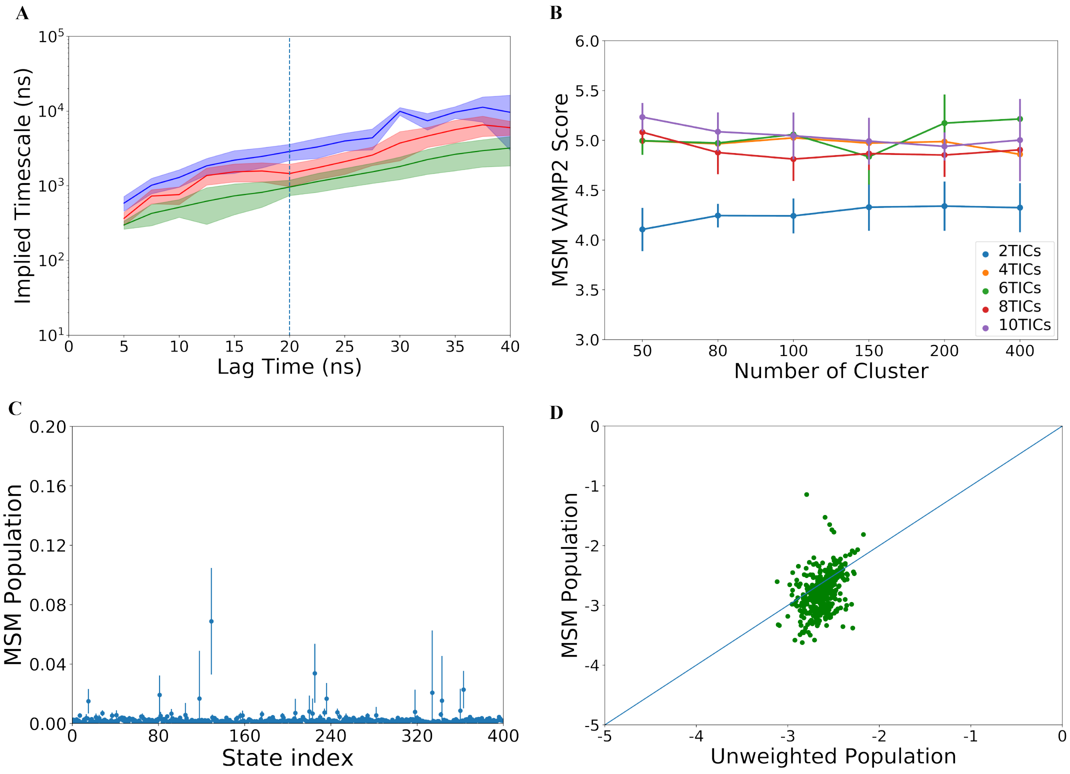


Figure S10: **Markov state model construction and analysis for dehydrated ProcA3.3 WT.** (A) Implied timescale calculated from the 2nd (blue), 3rd (red) and 4th (green) highest MSM eigenvalues at different lag times, MSM lagtime of 20 ns was chosen. (B) VAMP-2 scores with different clusters and different number of tICs of MSM, 400 clusters and 6 tic dimensions were selected for MSM. (C) MSM population density of different MSM state index, error bar was calculated by 200 rounds bootstrapping with 80% of total trajectories. (D) MSM population vs Raw population in each state.

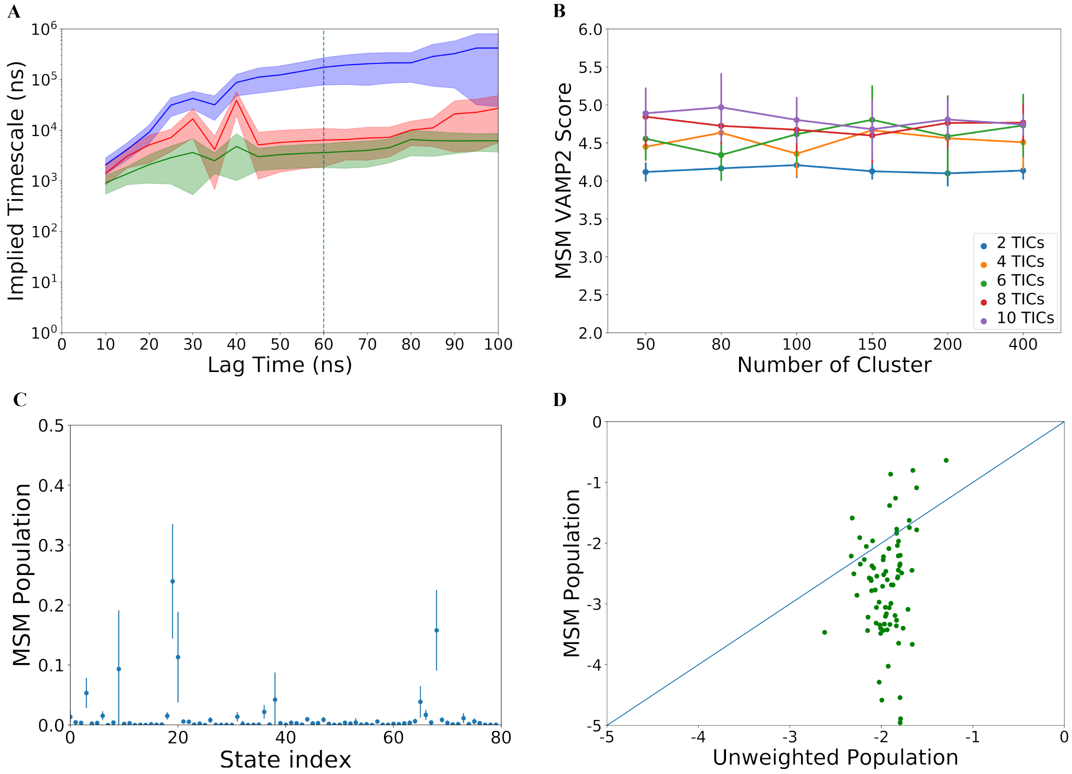


Figure S11: **Markov state model construction and analysis for dehydrated ProcA3.3 Variant1.** (A) Implied timescale calculated from the 2nd (blue), 3rd (red) and 4th (green) highest MSM eigenvalues at different lag times, MSM lagtime of 60 ns was chosen. (B) VAMP-2 scores with different clusters and different number of tICs of MSM, 80 clusters and 10 tic dimensions were selected for MSM. (C) MSM population density of different MSM state index, error bar was calculated by 200 rounds bootstrapping with 80% of total trajectories. (D) MSM population vs Raw population in each state.

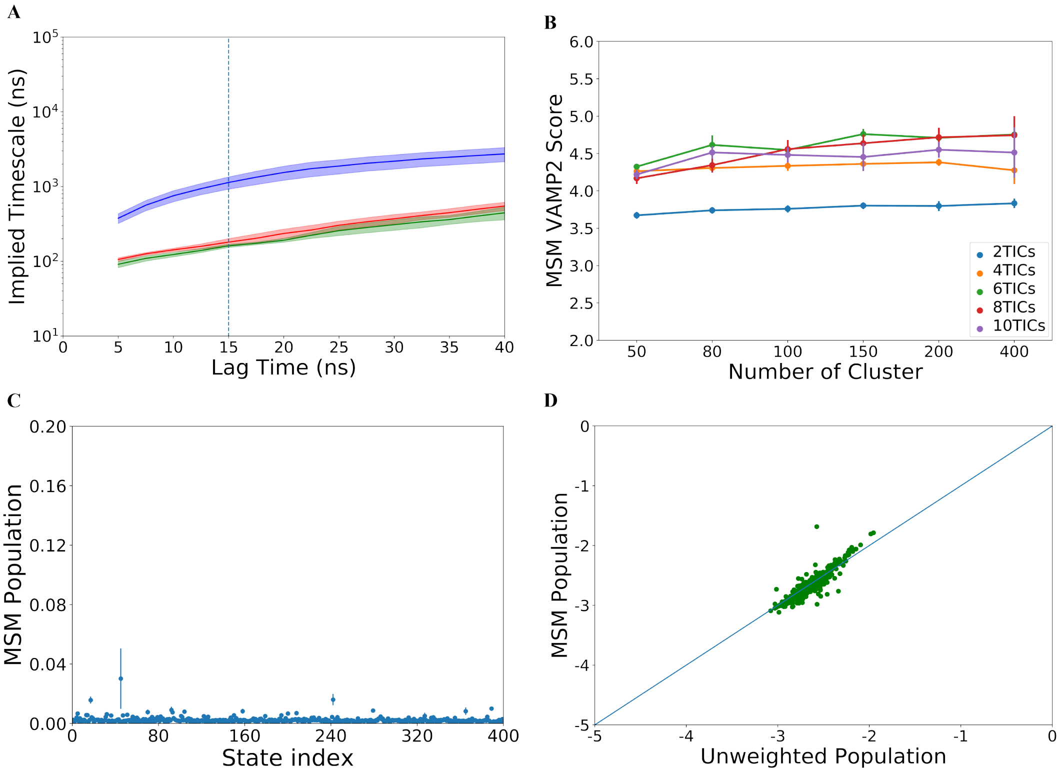


Figure S12: **Markov state model construction and analysis for ProcA1.1.** (A) Implied timescale calculated from the 2nd (blue), 3rd (red) and 4th (green) highest MSM eigenvalues at different lag times, MSM lagtime of 15 ns was chosen. (B) VAMP-2 scores with different clusters and different number of tICs of MSM, 400 clusters and 8 tic dimensions were selected for MSM. (C) MSM population density of different MSM state index, error bar was calculated by 200 rounds bootstrapping with 80% of total trajectories. (D) MSM population vs Raw population in each state.

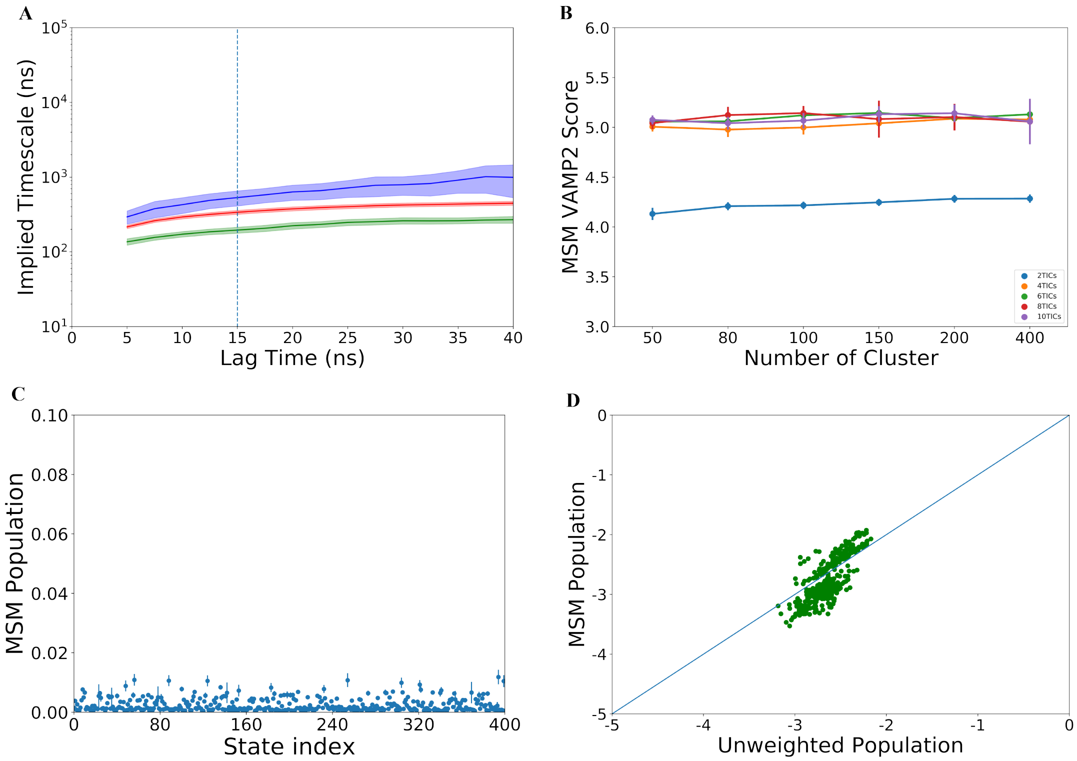


Figure S13: **Markov state model construction and analysis for ProcA2.8.** (A) Implied timescale calculated from the 2nd (blue), 3rd (red) and 4th (green) highest MSM eigenvalues at different lag times, MSM lagtime of 15 ns was chosen. (B) VAMP-2 scores with different clusters and different number of tICs of MSM, 400 clusters and 6 tic dimensions were selected for MSM. (C) MSM population density of different MSM state index, error bar was calculated by 200 rounds bootstrapping with 80% of total trajectories. (D) MSM population vs Raw population in each state.

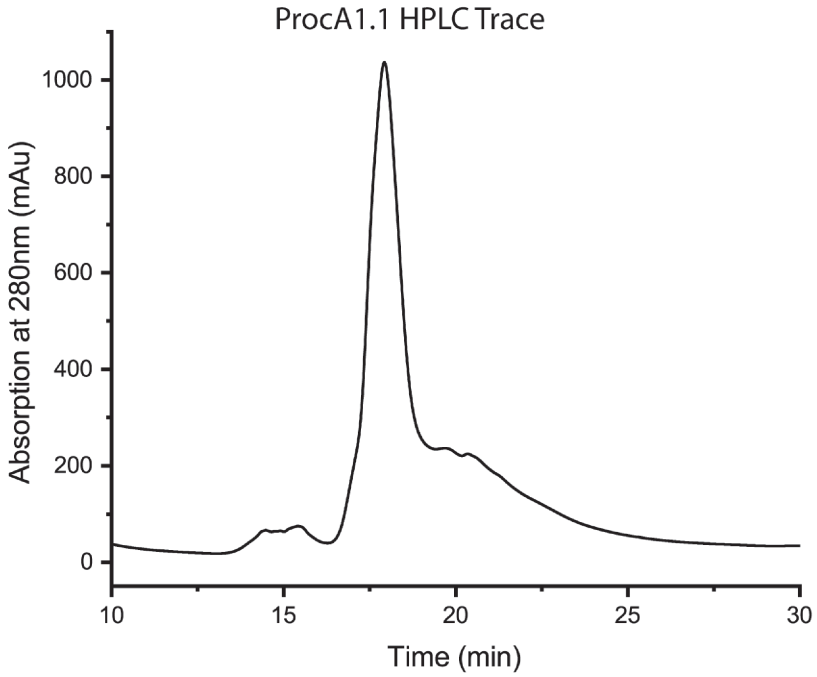


Figure S14: Chromatogram of ProcA1.1 purified by high-performance liquid chromatography. ProcA1.1 elutes with a retention time of 18 minutes.

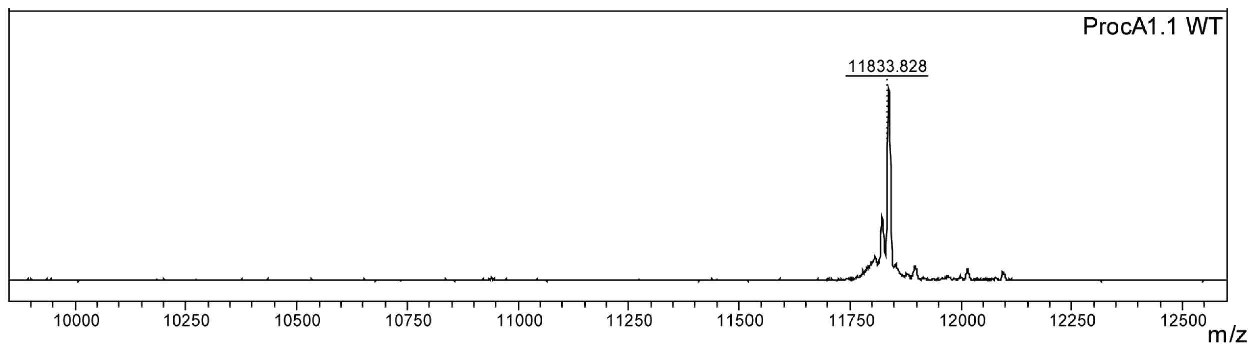


Figure S15: MALDI-TOF mass spectrum of HPLC purified unmodified ProcA1.1 (calculated m/z= 11833.832 ;observed= 11833.828).

Table S1: Molecular Dynamic Simulation Systems

| Peptide | Status | Sequence | Simulation Time |
|---|---|---|---|
| ProcA3.3 WildType | Unmodified | GDTGIQAVLHTAGCYGGTKMCRA | 40$\mu$s |
| ProcA3.3 WildType | Dehydrated | GD**Dhb**GIQAVLH**Dhb**AGCYGG**Dhb**KMCRA | 25$\mu$s |
| ProcA3.3 Variant1 | Unmodified | GDTGIQAVLHTLICFVITHLCRA | 40$\mu$s |
| ProcA3.3 Variant1 | Dehydrated | GD**Dhb**GIQAVLH**Dhb**LICFVI**Dhb**HLCRA | 25$\mu$s |
| ProcA3.3 Variant2 | Unmodified | GDTGIQAVLHTLFCYGGTFDCRA | 40$\mu$s |
| ProcA1.1 | Unmodified | FFCVQGTANRFTINVC | 25$\mu$s |
| ProcA2.8 | Unmodified | AACHNHAPSMPPSYWEGEC | 25$\mu$s |

Residues in red are residues varied in ProcA3.3 variants.
